# Changes in dorsomedial striatum activity mediate expression of goal-directed vs. habit-like cue-induced cocaine seeking

**DOI:** 10.1101/2023.07.24.550364

**Authors:** Brooke N. Bender, Sierra J. Stringfield, Mary M. Torregrossa

**Affiliations:** Department of Psychiatry, University of Pittsburgh, 450 Technology Drive, Pittsburgh, PA 15219, United States; Center for Neuroscience, University of Pittsburgh, 4200 Fifth Ave, Pittsburgh, PA 15213, United States

## Abstract

A preclinical model of cue exposure therapy, cue extinction, reduces cue-induced cocaine seeking when drug seeking is goal-directed but not habitual. Goal-directed and habitual behaviors differentially rely on the dorsomedial striatum (DMS) and dorsolateral striatum (DLS), but the effects of cue extinction on dorsal striatal responses to cue-induced drug seeking are unknown. We used fiber photometry to examine how dorsal striatal intracellular calcium and extracellular dopamine activity differs between goal-directed and habitual cue-induced cocaine seeking and how it is impacted by cue extinction. Rats trained to self-administer cocaine paired with an audiovisual cue on schedules of reinforcement that promote goal-directed or habitual cocaine seeking had different patterns of dorsal striatal calcium and dopamine responses to cue-reinforced lever presses. Cue extinction reduced calcium and dopamine responses during subsequent drug seeking in the DMS, but not in the DLS. Therefore, cue extinction may reduce goal-directed behavior through its effects on the DMS, whereas habitual behavior and the DLS are unaffected.

## Introduction

A major obstacle in the treatment of substance use disorders (SUDs) is maladaptive learning and memory, which can promote drug craving, use, and relapse (*1*–*5*). Several types of associative learning contribute to persistent drug use, and drug exposure also enhances learning and the strength of these associative memories (*6*–*12*). Response-outcome learning occurs when a desired outcome, such as a drug of abuse, becomes associated with a behavioral response, or action, that produces the drug effect (*13*–*15*). Response-outcome associations can then promote goal-directed drug-seeking behavior (*16*). As learning continues and an action repeatedly leads to the same outcome, stimulus-response associations begin to form where the environmental stimuli present during response-outcome learning (e.g, contexts or discrete cues) become sufficient to drive the behavioral response independent of the value of the outcome, which is defined as habitual behavior (*13*–*15*). Therefore, over time these stimuli alone can promote putatively habitual drug-seeking behaviors (*15*, *17*). Finally, in addition to action-related learning, Pavlovian associations also form between environmental stimuli and the effects of a drug, which can also promote motivated behavior (*10*, *16*, *18*). The presentation of drug-associated stimuli, or cues, has been shown to enhance subjective levels of drug craving in individuals with SUDs, promote relapse, and activate implicated brain regions, including the nucleus accumbens, dorsal striatum, and regions of the cortex (*1*, *4*, *5*, *19*–*24*).

Therefore, the extinction of Pavlovian drug-cue associations has been proposed as a potential therapeutic target in the treatment of SUDs (*25*–*27*). Cue exposure therapy, the repeated presentation of cues in the absence of the associated outcome, has been shown to be an effective behavioral treatment for others psychiatric disorders that involve maladaptive Pavlovian associations, such as phobias and post-traumatic stress disorder (*28*, *29*). Additionally, cue extinction, a preclinical model of cue exposure therapy, reduces cue-induced cocaine seeking in rodent cocaine self-administration models (*30*–*33*). However, the clinical application of cue exposure therapy to SUDs has yielded modest results (*26*, *34*). There are likely several reasons for this difficulty in translation, including context dependency (*35*–*37*). However, our lab has also shown that Pavlovian cue extinction reduces goal-directed cocaine seeking, but has no effect on habitual cocaine seeking unless goal-directed control is restored (*38*). Therefore, a lack of effect of cue extinction on habitual components of drug seeking may also be a contributing factor to this difficulty in translation.

Extensive literature has implicated distinct neural circuits in goal-directed and habitual behavior (*18*, *39*, *40*). Dopaminergic inputs from the substantia nigra to the dorsomedial striatum (DMS) and dorsolateral striatum (DLS) are important for the initiation of goal-directed and habitual behavior, respectively (*41*–*44*). Additionally, other direct and indirect inputs to the dorsal striatum, including those from the cortex, thalamus, and amygdala, may be important for toggling between reliance on goal-directed and habitual behavior (*45*–*50*). Dopamine release in the DMS and DLS during operant reward seeking can differ between regions depending on the operant task and extent of training (*51*–*55*). Several studies using *in vivo* electrophysiology to compare DMS and DLS activity during operant reward-seeking behavior have also shown distinct patterns of neural activity in these regions and indicate these patterns change as habitual behavior develops (*56*–*59*). However, the specific contributions of dopaminergic and other inputs to the dorsal striatum’s response to drug-associated cues and how they might be impacted by Pavlovian cue extinction remain unclear.

In the present study, we employed fixed-ratio (FR) and second-order (SO) schedules of reinforcement to facilitate either goal-directed (DMS dopamine-dependent) or habitual (DLS dopamine-dependent) cocaine seeking, respectively, as previously described (*38*, *42*, *50*). We utilized fiber photometry to examine dorsal striatal calcium and dopamine activity during drug seeking throughout the establishment of goal-directed and habitual cocaine self-administration and evaluated the effects of cue extinction on activity in these regions. We found distinct signatures of calcium and dopamine activity in the dorsal striatum in rats trained on FR or SO reinforcement schedules to promote goal-directed or habitual cocaine seeking, respectively. Additionally, we showed that cue extinction impacted DMS, but not DLS, calcium and dopamine activity during subsequent drug seeking, which suggests that cue extinction does not impact the neural circuitry promoting habitual behavior.

## Results

### FR- and SO-trained rats do not differ in daily cocaine self-administration or cue-induced drug-seeking after cue extinction

The dopamine fluorescent sensor dLight and calcium sensor RCaMP were expressed contralaterally in the aDLS and pDMS in male and female rats (n=26), and optic fibers and jugular vein catheters were implanted (Figure 1A). After all experiments, virus expression at the base of the fiber (Figure 1B) and fiber placement (Figure 1C) were confirmed by fluorescent microscopy. A total of 10 rats were excluded due to death after surgery, loss of catheter patency, or bilateral fiber misplacement or virus expression. Remaining rats (n=16; 9 males and 7 females) were trained to self-administer cocaine (1 mg/kg/inf) for 20 days before they underwent cue extinction and a subsequent cue-induced drug-seeking test (Figure 1D). During daily self-administration, there was a main effect of training day on number of infusions (F_(4.776,66.86)_=5.759, p=0.0002), but no main effect of training schedule (F_(1,14)_=0.002179, p=0.9634) or interaction (F_(19,266)_=1.486, p=0.0899) (2-way rmANOVA) (Figure 1E). For lever presses, there was a 3-way training day × schedule × lever interaction (F_(19,266)_=9.801, p<0.0001) (3-way rmANOVA) (Figure1F). These data indicate that both groups self-administered more cocaine infusions and made more active lever presses as training progressed, and that the increase in active lever presses was more pronounced in SO-trained rats, which is expected because these rats had to increase their lever presses to receive the same number of infusions. During the cue-induced drug-seeking test after cue extinction, there was no difference in the ratio of active lever presses during test to the final day of self-administration between FR-trained and SO-trained rats (p=0.3229, η^2^=0.08873) (unpaired *t*-test) (Figure 1G).

**Fig. 1.**
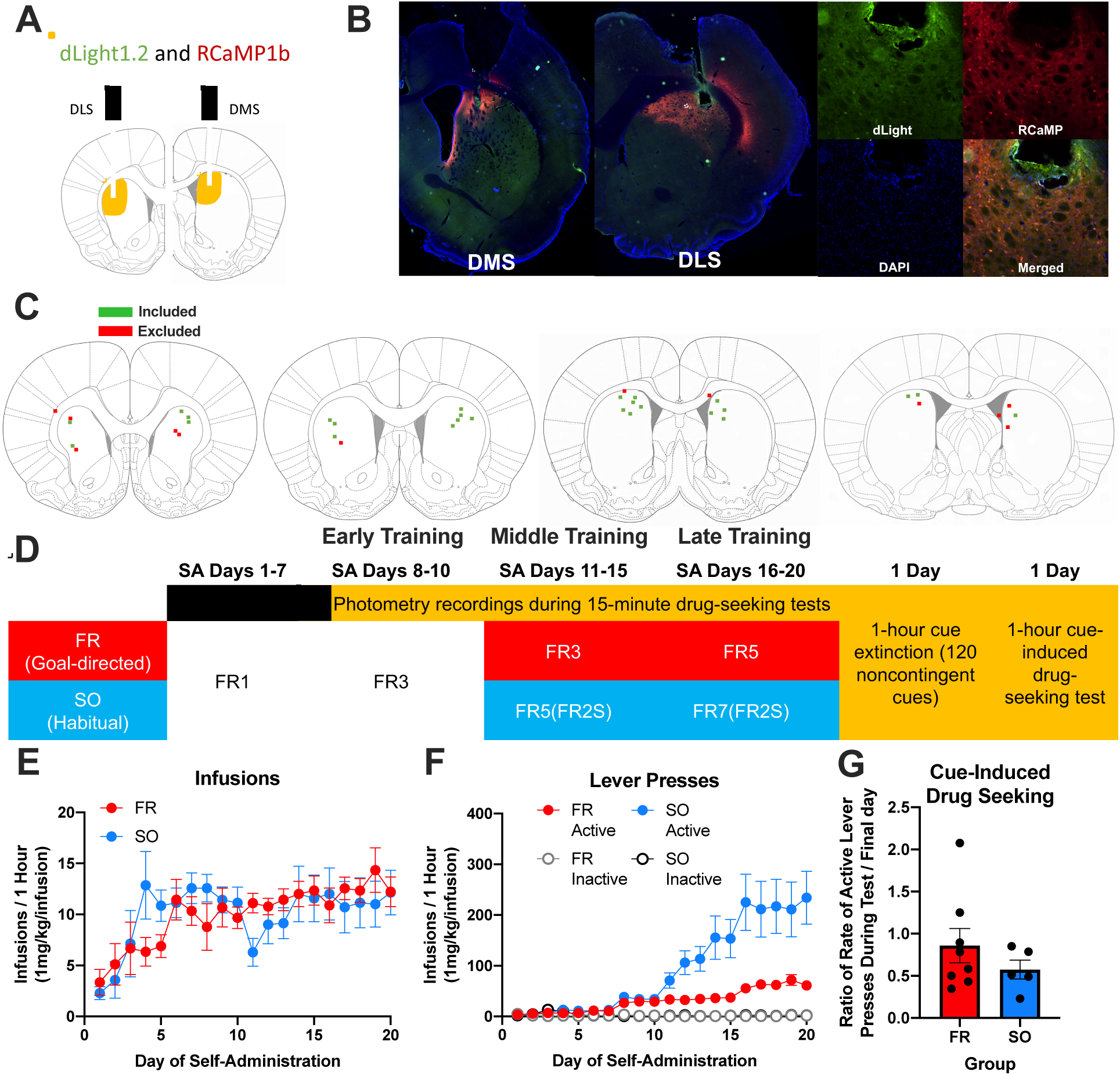
FR- and SO-trained rats do not differ in daily cocaine self-administration or cue-induced drug-seeking after cue extinction. Schematic for fiber placement and virus expression in the DLS and DMS (**A**). Representative images of fiber placement and virus expression in the DMS and DLS (left) and at higher magnification in the DLS (right) with fluorescent channels shown individually and merged and fiber locations outlined in dotted white lines (**B**). For all rats, fiber placement and virus expression were evaluated via fluorescent microscopy, and green bars represent fibers appropriately placed with confirmed virus expression at the base of the fiber, while red bars indicate fibers excluded from analysis due to either fiber misplacement and/or lack of virus expression (**C**). Rats were trained to self-administer cocaine for 20 days on different schedules of reinforcement (FR or SO) before undergoing cue extinction and a subsequent cue-induced drug-seeking test, and after acquisition, photometry recordings occurred in 15-minute drug-seeking tests that preceded daily self-administration, during cue extinction, and during the subsequent cue-induced drug-seeking test (**D**). During self-administration, there was a main effect of training day on the number of daily cocaine infusions, where infusions increased for both groups as training progressed (**E**). For lever presses during daily self-administration, there was a 3-way training day × training schedule × lever interaction (**F**). There was no difference between groups in the ratio of active lever presses during the post-cue extinction cue-induced drug-seeking test compared to the final day of self-administration (**G**). Graphs show group means ± SEM and individual data points where possible.

### After acquisition, dorsal striatal calcium responses are greater for cue-reinforced than unreinforced active lever presses

After 7 days of cocaine self-administration on an FR1 schedule, all rats self-administered on an FR3 schedule for 3 days. Following the first day of FR3 training, fiber photometry recordings took place in daily 15-minute drug-seeking tests prior to self-administration. The first 3 days of recording to examine dorsal striatal responses during “early training” occurred after acquisition but before rats were split into FR- and SO-trained groups. Because we wanted to determine if there were any differences between rats later split into groups, during this early training phase we first compared dorsal striatal responses between rats that would later be separated into FR- and SO-trained groups. On the schedules of reinforcement used in these studies, some active lever presses resulted in cue presentation upon the completion of the reinforcement schedule, but others that occurred before schedule completion had no consequences. Because we were particularly interested in the role of the drug-paired cue, we compared calcium and dopamine responses in the dorsal striatum between future training groups for cue-reinforced versus unreinforced active lever presses using 2-way ANOVAs. Therefore, any differences were due to differences in the response to cue presentation after lever press and were not a motion artifact of performing the lever press. Because we used both male and female rats, we first determined that there was no effect of sex or estrous phase on dorsal striatal responses to lever presses (Figure S1). Therefore, males and females were combined for analyses throughout.

During early training prior to splitting animals into groups, there was a main effect of cue reinforcement (F_(1,8)_=12.98, p=0.0070) on calcium peak z-score amplitude in the DLS in the 1 second after lever press, but no effect of future training schedule (F_(1,8)_=4.263, p=0.0728) or interaction (F_(1,8)_=2.214, p=0.1751) (Figure 2A). Similarly, for calcium peak amplitude in the DMS, there was a main effect of cue reinforcement (F_(1,9)_=9.035, p=0.0148), but no effect of future training schedule (F_(1,9)_=0.006144, p=0.9392) or interaction (F_(1,9)_=0.1558, p=0.7022) (Figure 2B). For dopamine peak amplitude in the DLS, there were no main effects of cue reinforcement (F_(1,9)_=0.06979, p=0.7976) or future training schedule (F_(1,9)_=1.907, p=0.2006) or interaction (F_(1,9)_=0.2988, p=0.5979) (Figure 2C). Similarly, for dopamine peak amplitude in the DLS, there were no main effects of cue reinforcement (F_(1,9)_=0.06176, p=0.8093), future training schedule (F_(1,9)_=0.0007846, p=0.9783) or interaction (F_(1,9)_=0.02546, p=0.8767) (Figure 2D). Overall, these data suggest that after acquisition of cocaine self-administration, calcium activity in the DMS and DLS was greater in response to cue-reinforced lever presses compared to lever presses that did not result in cue presentation. Interestingly, cue reinforcement after lever press did not impact dopamine activity in either region during this early training phase. Throughout, we present peak amplitude data in the main figures, but we also calculated the area under the curve (AUC), which can be found in supplemental figures S1-S6. During early training, results from AUC data were similar to peak data, and there were no significant correlations between the average number of active lever presses during drug-seeking tests and the average calcium or dopamine peak amplitude or AUC for cue-reinforced lever presses (Figure S2), which suggests that different rates of lever pressing do not impact dorsal striatal responses to cues.

**Fig. 2.**
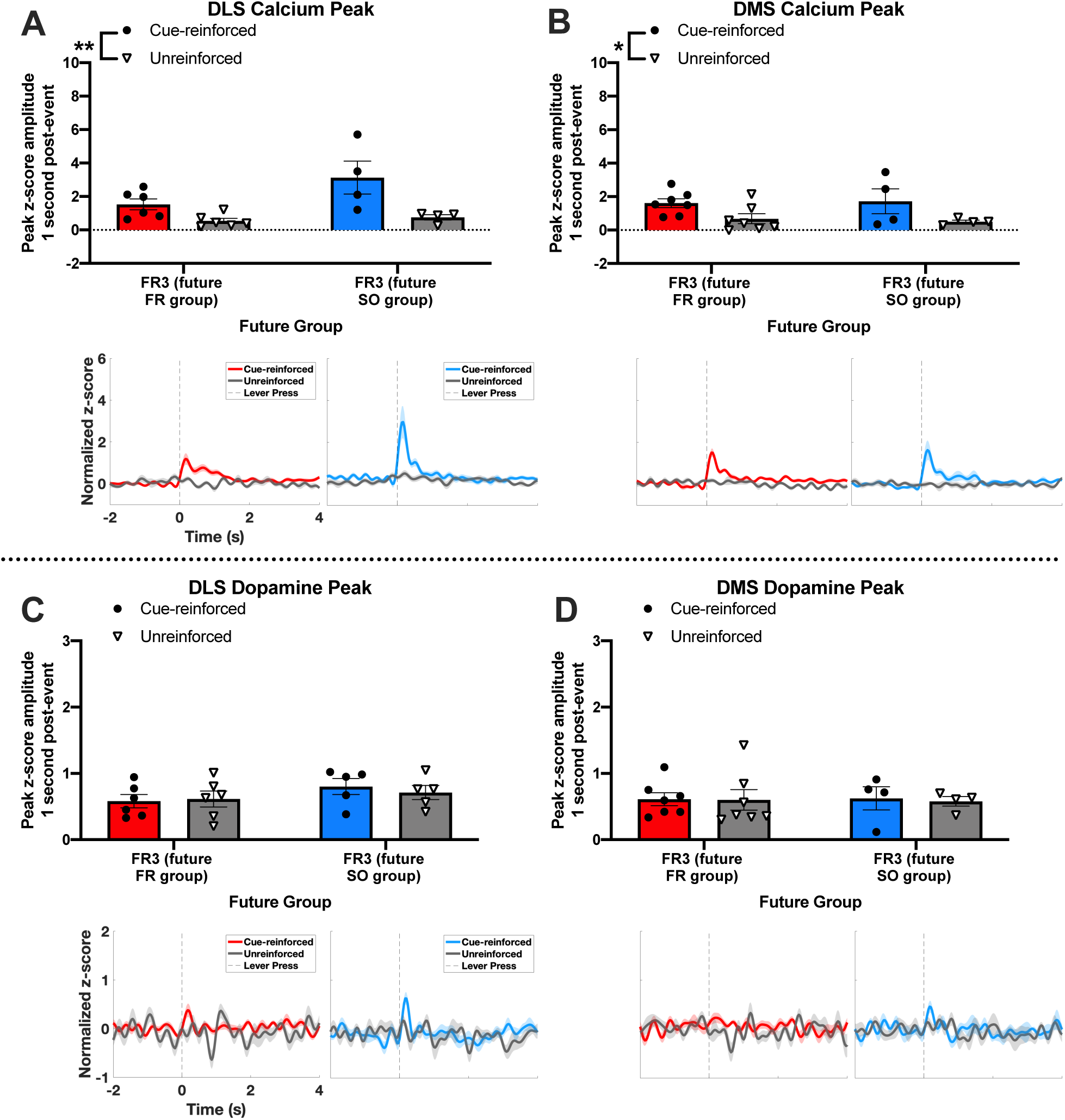
After acquisition, dorsal striatal calcium responses are greater for cue-reinforced than unreinforced active lever presses. After acquisition and prior to splitting rats into FR- and SO-trained groups, fiber photometry recordings occurred during 15-minute drug-seeking tests prior to daily self-administration, during which some active lever presses had no consequence (unreinforced) and an active lever press that completed the FR3 schedule resulted in cue presentation (cue-reinforced) and timeout (levers retracted, houselight extinguished). For DLS (**A**) DMS (**B**) calcium, there was a main effect of cue reinforcement on peak z-score amplitude in the 1 second after lever press, but there was no effect of future training schedule or interaction. For DLS (**C**) and DMS (**D**) dopamine peak z-score amplitude, there was no effect of cue reinforcement, future training group, or interaction. Graphs show group means ± SEM and individual data points. Traces show overall average trace for each event for each future group aligned to behavioral events with SEM shown with shading and dashed vertical lines indicating time of lever press. *p<0.05; **p<0.01.

### FR-trained rats, but not SO-trained rats, show greater DMS calcium activity after cue-reinforced compared to unreinforced lever presses

After 10 days of self-administration, rats were split into FR-trained and SO-trained groups and trained on different schedules of reinforcement accordingly. The next 10 days were split into 5 days of middle training and 5 days of late training. During SO-schedule training, short 1-second cues are presented in addition to 20-second cues presented upon schedule completion, and for some measures SO-trained rats had different responses to short cues compared to long cues (Figure S3). Therefore, only long cues were compared between FR- and SO-trained rats. To determine if SO-trained rats had different dorsal striatal responses to lever presses than FR-trained rats, we compared cue-reinforced to unreinforced lever presses between groups for each phase of training (middle or late) using 3-way ANOVAs. For DLS calcium peak amplitude, there was a main effect of cue reinforcement (F_(1,9)_=31.21, p=0.0003), but no main effects of training schedule (F_(1,9)_=4.259, p=0.0691) or phase of training (F_(1,9)_=3.205, p=0.1633), and there were no cue reinforcement × training schedule (F_(1,9)_=0.00748, p=0.9318), cue reinforcement × phase of training (F_(1,9)_=0.6451, p=0.4426), phase of training × training schedule (F_(1,9)_=3.954, p=0.0780), or 3-way interactions (F_(1,9)_=0.6999, p=0.4245) (Figure 3A). For DMS calcium peak amplitude, there was a main effect of cue reinforcement (F_(1,9)_=9.569, p=0.0129) and a cue reinforcement × training schedule interaction (F_(1,9)_=5.370, p=0.0457), but no main effect of training schedule (F_(1,9)_=1.870, p=0.2047) or phase of training (F_(1,9)_=1.130, p=0.3155), and no cue reinforcement × phase of training (F_(1,9)_=0.003176, p=0.9563), phase of training × training schedule (F_(1,9)_=4.718, p=0.0579), or 3-way interaction (F_(1,9)_=0.01580, p=0.9027) (Figure 3B). These data suggest that while both FR-trained and SO-trained rats had greater calcium responses in the DLS to cue-reinforced than unreinforced lever presses, this difference was only present in the DMS for FR-trained rats, but not in SO-trained rats. In other words, SO training led to a loss of cue-induced calcium-indicated activity selectively in the DMS.

**Fig. 3.**
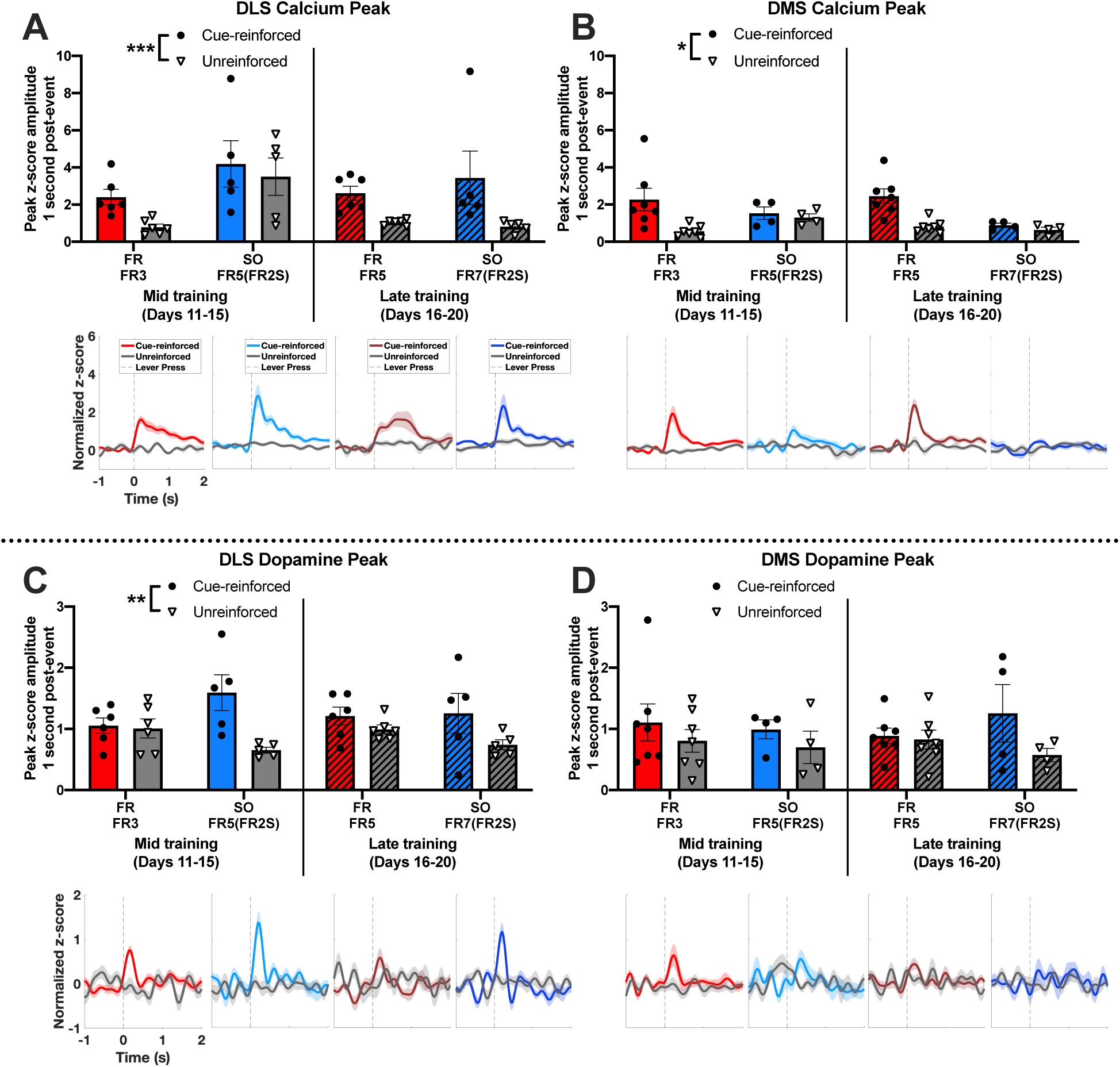
FR- and SO-trained rats have different patterns of dorsal striatal calcium and dopamine activity during drug seeking. Rats were separated into FR-trained and SO-trained groups for the remaining 10 days of self-administration and trained on different schedules of reinforcement accordingly for the middle and late phases of training. Dorsal striatal calcium and dopamine responses to cue-reinforced and unreinforced lever presses were compared for each training schedule and phase of training. There was a main effect of cue reinforcement on DLS calcium peak amplitude, but no main effects of training schedule or phase of training or interactions (**A**). For DMS calcium peak amplitude, there was a main effect of cue reinforcement and a cue reinforcement × training schedule interaction, but no other main effects or interactions (**B**). There was a main effect of cue reinforcement and a cue reinforcement × training schedule interaction for DLS dopamine peak amplitude (**C**), but there were no main effects or interactions for DMS dopamine peak amplitude (**D**). Graphs show group means ± SEM and individual data points. Traces show overall average trace for each event for each group aligned to behavioral events with SEM shown with shading and dashed vertical lines indicating time of lever press. *p<0.05; **p<0.01; ***p<0.001.

### SO-trained rats, but not FR-trained rats, show greater DLS dopamine responses to cue-reinforced compared to unreinforced lever presses

For DLS dopamine peak amplitude during middle and late training, there was a main effect of cue reinforcement (F_(1,9)_=11.42, p=0.0081) and a cue reinforcement × training schedule interaction (F_(1,9)_=5.494, p=0.0437), but no main effect of training schedule (F_(1,9)_=0.001535, p=0.9696) or phase of training (F_(1,9)_=0.06291, p=0.8076), and no cue reinforcement × phase of training (F_(1,9)_=0.3683, p=0.5589), phase of training × training schedule (F_(1,9)_=0.8370, p=0.3841), or 3-way interaction (F_(1,9)_=1.951, p=0.1960) (Figure 3C). There were no main effects of cue reinforcement (F_(1,9)_=0.2947, p=0.1202), training schedule (F_(1,9)_=0.01592, p=0.9024), or phase of training (F_(1,9)_=0.008296, p=0.9294) on DMS dopamine peak amplitude, and there were no cue reinforcement × training schedule (F_(1,9)_=6271, p=0.4488), cue reinforcement × phase of training (F_(1,9)_=0.1141, p=0.7433), phase of training × training schedule (F_(1,9)_=0.2727, p=0.6141), or 3-way interactions (F_(1,9)_=2.060, p=0.1851) (Figure 3D). These data suggest that SO-trained rats, but not FR-trained rats, had increased DLS dopamine activity after cue-reinforced lever presses, but neither group had greater DMS dopamine in response to cue-reinforced compared to unreinforced lever presses. Data for AUC of calcium and dopamine responses during middle and late training are presented in the supplement (Figure S3).

### Cue extinction reduces DMS calcium peak amplitude during drug seeking selectively in FR-trained rats

Given that we have previously found that SO-trained rats are resistant to the effects of cue extinction in modulating their cocaine-seeking behavior, and that this is reliant on activity in the DLS (*38*), we wanted to determine if cue extinction differentially affected dorsal striatal activity in a drug seeking test after cue extinction. Rats underwent a 1-hour cue-induced drug-seeking test during which photometry recordings occurred. Dorsal striatal calcium and dopamine peak amplitudes after cue-reinforced or unreinforced lever presses during this post-cue extinction drug-seeking test (post-ext) were compared to the late phase of training (pre-ext) using 3-way ANOVAs. There was a main effect of cue reinforcement (F_(1,7)_=8.302, p=0.0236) on DLS calcium peak amplitude, but no main effects of training schedule (F_(1,7)_=3.393, p=0.2080) or cue extinction (F_(1,7)_=0.4329, p=0.5316) or cue reinforcement × training schedule (F_(1,7)_=1.480, p=0.2632), cue reinforcement × cue extinction (F_(1,7)_=0.06984, p=0.7992), training schedule × cue extinction (F_(1,7)_=1.385, p=0.2778), or 3-way interactions (F_(1,7)_=0.3617, p=0.5665) (Figure 4A). These data suggest that cue extinction did not impact DLS calcium activity for either FR- or SO-trained rats. For DMS calcium peak amplitude, there was a main effect of cue reinforcement (F_(1,7)_=11.42, p=0.0118) and a main effect of cue extinction (F_(1,7)_=6.514, p=0.0380), as well as significant cue reinforcement × training schedule (F_(1,7)_=6.362, p=0.0397) and training schedule × cue extinction interactions (F_(1,7)_=7.953, p=0.0258) (Figure 4B). There was no main effect of training schedule (F_(1,7)_=2.980, p=0.1279) on DMS dopamine peak amplitude, and there was no cue reinforcement × cue extinction (F_(1,7)_=0.1234, p=0.7358) or 3-way interaction (F_(1,7)_=0.04091, p=0.8455) (Figure 4B). These results suggest that cue extinction resulted in a reduction in DMS calcium peak amplitude for FR-trained rats, while SO-trained rats were unaffected, which may be due to their already minimal DMS calcium response. Effects of cue extinction on calcium AUC can be found in the supplement (Figure S4).

**Fig. 4.**
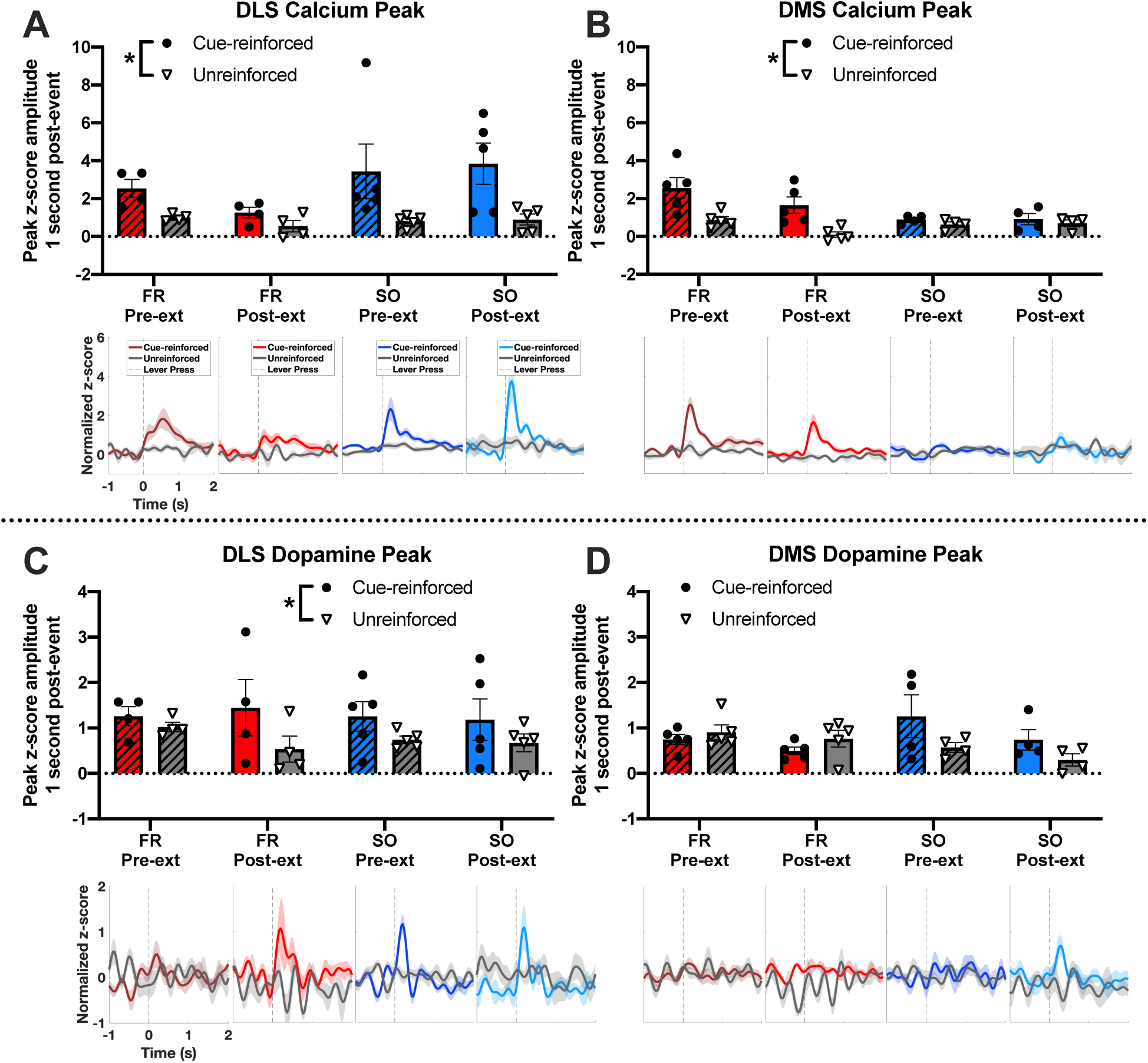
Cue extinction results in changes in DMS, but not DLS, calcium and dopamine activity during drug seeking. To examine the effects of cue extinction on dorsal striatal calcium and dopamine activity, peak amplitudes during the post-cue extinction drug-seeking test (post-ext) were compared to the late phase of training (pre-ext). There was a main effect of cue reinforcement on DLS calcium peak amplitude, but no effects of cue extinction or training schedule or interactions (**A**). For DMS calcium peak amplitude, there was a main effect of cue reinforcement, a main effect of cue extinction, and significant cue reinforcement × training schedule and training schedule × cue extinction interactions, but no other interactions (**B**). There was a main effect of cue reinforcement on DLS dopamine peak amplitude, with no other main effects or interactions (**C**). Finally, there was a main effect of cue extinction on DMS dopamine peak amplitude, but no other effects or interactions (**D**). Graphs show group means ± SEM and individual data points. Traces show overall average trace for each event for each group aligned to behavioral events with SEM shown with shading and dashed vertical lines indicating time of lever press. *p<0.05.

### Cue extinction reduces DMS dopamine peak amplitudes during drug seeking in both groups

For DLS dopamine peak amplitude, there was a main effect of cue reinforcement (F_(1,7)_=6.724, p=0.0358), but no effect of training schedule (F_(1,7)_=0.1056, p=0.7547) or cue extinction (F_(1,7)_=0.2119, p=0.6592), and there were no cue reinforcement × training schedule (F_(1,7)_=0.02193, p=0.8865), cue reinforcement × cue extinction (F_(1,7)_=2.090, p=0.1915), training schedule × cue extinction (F_(1,7)_=0.02783, p=0.8722), or 3-way interactions (F_(1,7)_=2.151, p=0.1859) (Figure 4C). There was a main effect of cue extinction (F_(1,7)_=8.889, p=0.0205) on DMS dopamine peak amplitude, but there was no main effect of cue reinforcement (F_(1,7)_=1.106, p=0.3280) or training schedule (F_(1,7)_=0.004638, p=0.9476), and there were no cue reinforcement × training schedule (F_(1,7)_=5.388, p=0.0533), cue reinforcement × cue extinction (F_(1,7)_=0.6134, p=0.4592), training schedule × cue extinction (F_(1,7)_=1.005, p=0.3495), or 3-way interactions (F_(1,7)_=0.07993, p=0.7856) (Figure 4D). These results suggest that cue extinction resulted in an overall reduction in DMS dopamine peak amplitude after any lever press, regardless of cue reinforcement, in both groups, while DLS dopamine responses were not affected. Effects of cue extinction on dopamine AUC can be found in the supplement (Figure S4).

## Discussion

In the present study, we show distinct patterns of calcium and dopamine activity during drug seeking in the dorsal striatum in rats trained on different schedules of reinforcement, where SO training results in an enhanced DLS dopamine response and a reduced DMS calcium response to cue-reinforced lever presses compared to FR training. Additionally, we show evidence that cue extinction impacts DMS, but not DLS, calcium and dopamine activity during later drug-seeking, which suggests that extinction of the Pavlovian cocaine-cue association impacts the DMS circuitry important for goal-directed drug seeking, but does not impact the DLS circuitry important for habitual drug seeking (*42*, *50*). We have previously shown that cue extinction does not affect cue-induced drug seeking in rats trained on SO schedules of reinforcement to promote DLS dopamine-dependent behavior unless goal-directed behavior is restored (*38*). In the present study, cue extinction’s lack of effect on the DLS provides a biological explanation for why cue extinction does not affect habitual drug seeking. These findings add to existing literature that indicates divergent roles of the DMS and DLS in goal-directed and habitual drug seeking and expand our understanding of how cocaine-cue associations facilitate dorsal striatal activity. In these experiments, we chose to use fiber photometry, which allows for the *in vivo* comparison of bulk changes in fluorescent output of fluorescent indicators in the regions of interest. Fluorescent dopamine sensors provide the advantage of enhanced temporal resolution compared to other methods of monitoring dopamine release *in vivo*, including microdialysis and fast-scanning cyclic voltammetry (FSCV) (*60*–*62*). Using fiber photometry also allowed us to simultaneously monitor dopamine release (via dLight) and intracellular calcium (via RCaMP) in the same rats (*63*).

Although *in vivo* electrophysiology has enhanced temporal sensitivity compared to fiber photometry, fiber photometry is more stable for long-term comparison across days, which was important for the present study (*63*). Additionally, monitoring intracellular calcium may provide distinct information compared to *in vivo* electrophysiology. It was initially proposed that intracellular calcium activity reported by calcium sensors like GCaMP and RCaMP are proxy indicators of neural activity, or cell firing (*64*). However, recent evidence suggests that at least in the dorsal striatum, where neurons have extensive dendritic arborization, changes in the fluorescent calcium signal reported by fiber photometry are more indicative of changes in non-somatic calcium and therefore do not reflect the same results as *in vivo* electrophysiology (*65*). Importantly, this evidence suggests that the changes in calcium fluorescence we report here may be interpreted as the summation of excitatory and inhibitory input into dorsal striatal neurons (*65*).

We initially recorded dorsal striatal dopamine and calcium activity during cue-induced drug seeking after acquisition. Notably, we did not show any significant differences in dopamine or calcium peak amplitudes between rats that would later be separated into FR- and SO-trained groups, which suggests that later differences in calcium and dopamine responses were the result of effects of training on different schedules of reinforcement and were not due to baseline differences in sensor expression. Additionally, our comparison between cue-reinforced and unreinforced active lever presses controls for motion artifacts that could be produced by the lever press action and isolates the contribution of cocaine-paired cue presentation on dorsal striatal activity (*66*).

After just one week of cocaine self-administration, there was an enhanced calcium response to cue-reinforced lever presses in both the DMS and DLS. Because these recordings took place after limited self-administration training at a timepoint when behavior would presumably be dependent on the DMS (*39*, *42*), the DLS calcium response to reinforced lever presses was somewhat surprising. There is some uncertainty about when the stimulus-response associations that later guide habitual behavior are learned. Typically, habitual behavior can be differentiated from goal-directed behavior through its lack of sensitivity to outcome devaluation and outcome contingency degradation (*15*). There is evidence that even after minimal operant training, DMS inhibition, DMS lesions, or exposure to stimulants can render behavior insensitive to outcome devaluation or contingency degradation (*7*, *8*, *39*, *67*, *68*). Insensitivity to these paradigms could suggest reliance on habitual behavior, but in some cases it could also be attributed to impaired execution of goal-directed behavior (*69*). Our results showing increased calcium activity after cue-reinforced lever presses in the DLS at this early timepoint, when behavior is presumably DMS-dependent and goal-directed, support the theory that stimulus-response learning occurs during early training, even though the behavioral response may still be goal-directed.

Interestingly, we did not show increased DMS dopamine release in response to cue-reinforced lever presses compared to unreinforced lever presses in either group at any phase of training. Because previous experiments have shown that after similar training, DMS dopamine antagonism reduces drug-seeking behavior, this finding was particularly surprising in the early training phase, as well as during the middle and late phase in FR-trained rats (*42*). The lack of effect is not likely due to limitations of the dopamine sensor dLight, as we did detect a DMS dopamine response to novel stimuli prior to training, in accordance with a previous study showing that midbrain dopamine neurons, including those that project to the dorsal striatum, respond to novel stimuli (*70*). It is well-established that dopamine release in the nucleus accumbens core occurs upon the presentation of a reward-predictive or reward-associated cue (*51*, *71*). However, the impact of reward-associated cues on DMS dopamine release is less clear. Because the DMS is particularly important for goal-directed drug seeking, which is promoted by the association between the lever press behavior and reward, DMS dopamine release may not be directly tied to cue presentation, but to reward delivery. Indeed, in mice that were trained to self-administer sucrose, calcium activity in dopamine neuron terminals in the DMS did show increased responses to nosepokes that were reinforced by sucrose delivery (*72*). Therefore, the data in the present study may not show a DMS dopamine response because we recorded DMS dopamine under extinction conditions, when lever presses were reinforced by the cue alone and not cocaine, due to technical limitations and to prevent cocaine exposure from impacting DMS dopamine. However, in rats trained to self-administer sucrose after a discriminative stimulus was presented, presentation of the discriminative stimulus resulted in increased DMS, but not DLS, dopamine release as measured by FSCV (*55*). In this case, the stimulus does not signal reward delivery, but the imminent availability of a lever that, when pressed, results in sucrose delivery (*55*). Therefore, future experiments should investigate the conditions for which DMS dopamine release occurs for reward-predictive or reward-associated cues. Interestingly, results from the present study did suggest DMS dopamine overall during drug seeking was reduced after cue extinction, even though this reduction was not specific to cue-reinforced lever presses.

On the other hand, our results indicate that SO, but not FR training, does enhance DLS dopamine release after cue-reinforced lever presses. These results agree with those of another study using in vivo microdialysis to measure dopamine release in the DLS in rats trained to self-administer cocaine on second-order schedules of reinforcement, which also showed enhanced DLS dopamine release during cue-induced drug seeking (*52*). Our findings complement these by providing enhanced temporal resolution and showing that this increased DLS dopamine release is specific to cue-reinforced lever presses. In another study, rats were trained to self-administer alcohol or sucrose on a variable-interval schedule, which usually promotes habitual behavior, and striatal dopamine release was measured with FSCV during operant responding for reward-associated cues (*15*, *51*). They observed enhanced dopamine release after cue-reinforced compared to unreinforced lever presses in the DLS, but not in the DMS (*51*). Our results extend these findings by indicating that SO-training to self-administer cocaine facilitates a similar pattern of dorsal striatal dopamine release, which suggests this enhanced DLS dopamine response to cue-reinforced lever presses is consistent across multiple reinforcers (cocaine, ethanol, and sucrose) and is also generalized between multiple schedules of reinforcement known to facilitate habitual behavior. Although FR-trained rats in the present study did not develop an enhanced DLS dopamine response to cue-reinforced lever presses, one previous study did report enhanced dopamine release in the DLS, measured by FSCV, after just 2-3 weeks of similar cocaine self-administration, but rats also received cocaine along with the cocaine-associated cue when they made an active nose poke (*54*). Therefore, reward delivery may also be required to promote this DLS dopamine response in FR-trained rats, and future experiments should determine what experimental parameters are required to facilitate a DLS dopamine response to cue-reinforced drug seeking.

With regard to calcium responses, we found increases in DMS responses to cue-reinforced compared to unreinforced lever presses through the late phases of training in FR-trained, but not SO-trained rats. Importantly, rats that would later be SO-trained did show increased DMS calcium responses during the early phase of training, which suggests that SO-training led to a loss of the DMS calcium response to cue-reinforced lever presses. Because calcium activity primarily reflects dendritic calcium, the reduction in calcium activity likely reflects reduced excitatory or enhanced inhibitory input into DMS neurons after SO training (*65*). Future studies should determine the contribution of different DMS inputs to the DMS calcium response to cue-reinforced behavior. Indeed, there is evidence that mPFC projections to the DMS are disengaged as a motor skill is refined, so it is possible that a reduction in excitatory mPFC input could occur in SO-trained rats (*73*). Additionally, the orbitofrontal cortex (OFC) is important for goal-directed behavior and modulates the DMS via both direct projections and indirectly through the other cortical areas and the amygdala, and could also be involved in this decrease in DMS calcium response in SO-trained rats (*45*, *74*). In addition to projections from the cortex and the amygdala, the DMS also receives thalamic inputs, which should also be investigated (*75*). Future studies could also evaluate the contribution of different neuronal sub-types, as direct- and indirect-pathway medium spiny neurons and the more sparse local interneurons may have different roles in these behaviors (*76*–*79*).

Next, we evaluated dorsal striatal calcium and dopamine responses to the non-contingent presentation of cocaine-paired cues during a Pavlovian cue extinction paradigm. Interestingly, DMS and DLS calcium responses to passive cue presentations were of a lower magnitude than those observed when the cue was presented after a lever press during drug seeking tests, and the response to cues did not change throughout the cue extinction session. Additionally, we did not observe a dorsal striatal dopamine response to noncontingent cues during cue extinction. These findings agree with those of another study showing that noncontingent cue presentations do not result in increased DLS dopamine release, as measured by *in vivo* microdialysis, after SO training to self-administer cocaine (*52*). Taken together, these results suggest that DLS dopamine release during habitual drug seeking is dependent upon both the lever press action as well as contingent cue presentation, further supporting the role of DLS dopamine in connecting cues (stimuli) with the lever press behavior.

Finally, we conducted an additional drug seeking test after cue extinction and compared dorsal striatal calcium activity to activity during the late phase of training, prior to cue extinction. In doing this comparison, we found that FR-trained rats had reduced DMS calcium responses to lever presses after cue extinction, but there was no effect in SO-trained rats or in either group in the DLS. These data suggest that the learned lack of association between cocaine and the drug-associated cue that occurs during cue extinction results in reduced DMS calcium activity during subsequent drug seeking, despite no change in the smaller DMS response to noncontingent cues throughout cue extinction. Additionally, the later reduction in DMS calcium activity after cue extinction learning selectively occurs in rats trained on a schedule that promotes goal-directed behavior. Inputs to the DMS that contribute to this reduction in DMS calcium activity after cue extinction are not currently known. Our lab has previously shown that cue extinction results in reduced synaptic strength of thalamo-amygdala synapses, and optogenetic depotentiation of these synapses mimics the effects of cue extinction (*32*). Therefore, it is possible that cue extinction affects DMS calcium activity by reducing input to the DMS either directly from the BLA or through the BLA’s interaction with the OFC, which could be the topic of future investigations (*45*, *74*). We have previously shown that cue extinction reduces cue-induced drug seeking in FR-trained, but not SO-trained rats (*38*). In our previous study, the effect of cue extinction in FR-trained rats was apparent when animals that underwent cue extinction were compared to those that did not. Therefore, the lack of difference in response ratio after cue extinction in FR-compared to SO-trained rats in the current study is consistent with our previous findings as we did not compare to a no extinction group (*38*). Interestingly, cue extinction affected DMS calcium activity in FR-trained rats and DMS dopamine activity in both groups, which suggests that cue extinction may inhibit circuitry that facilitates goal-directed action towards seeking cocaine under extinction conditions when cocaine is expected but not provided.

Overall, the present study expands upon previous literature examining the differential roles of the DMS and DLS in goal-directed and habitual behavior, respectively. We also present novel results showing that cue extinction reduces calcium and dopamine activity in the DMS during later drug seeking but has no effects on the DLS, which indicates that extinction of the cocaine-cue association impacts the circuitry involved in goal-directed, but not habitual cocaine seeking. Future experiments should examine specific projections to the DMS and how they are impacted by SO-schedule training and cue extinction. Together, these results provide novel insights into how cue extinction may reduce drug seeking that is goal-directed but not affect habitual drug seeking and indicate that future treatments for SUDs may need to address these aspects of drug-seeking behavior using different methods.

## Materials and Methods

### Experimental Design

Adult male and female rats expressing the fluorescent calcium indicator RCaMP1b and dopamine indicator dLight1.2 in the DMS and DLS were implanted with optic fibers in the DLS and DMS and jugular vein catheters. Rats were trained to self-administer cocaine paired with an audiovisual cue for 20 days, and were split into FR-trained and SO-trained groups and trained accordingly on different schedules of reinforcement to facilitate goal-directed (FR-trained) or habit-like (SO-trained) cocaine seeking behavior. Fiber photometry recordings under extinction conditions occurred during 15-minute drug-seeking tests that occurred immediately before self-administration on days 9-20 on the rat’s reinforcement schedule from the previous day. To determine how DLS and DMS calcium and dopamine responses are affected by lever presses that resulted in presentation of cocaine-associated cues, dorsal striatal responses were compared between active lever presses that resulted in cue presentation (cue-reinforced) and active lever presses that had no consequence (unreinforced). To examine how training schedule impacts dorsal striatal calcium and dopamine active, dorsal striatal responses were compared between FR-trained and SO-trained rats across early, middle, and late phases of training. Next, rats underwent fiber photometry recordings during cue extinction (120 20-second noncontingent cues) and a 1-hour cue-induced drug-seeking test. To determine if cue extinction impacts dorsal striatal calcium or dopamine activity, dorsal striatal responses were compared between the cue-induced drug-seeking test followed cue extinction and during the late phase of training.

### Animals

Adult Sprague-Dawley rats (Envigo) were 8-9 weeks old upon arrival (n=26; male n=14; female n=12). Animals were housed in auto-ventilated racks with automated watering in a temperature- and humidity-controlled room maintained on a 12-hour light-dark cycle. Rats were given ≥4 days to acclimate to the facility before surgical procedures and were pair-housed until catheter implantation. Rats had *ad libitum* access to food and water until 24 hours before the start of training, when they were food restricted to maintain ∼90% of their free-feeding body weight. Behavioral experiments were run in the light cycle and began within ∼3 hours of the same time each day. Procedures were conducted in accordance with the National Institute of Health’s *Guide for the Care and Use of Laboratory Animals* and were approved by the University of Pittsburgh’s Institutional Animal Care and Use Committee.

### Viral Vectors

Viral vectors encoding the fluorescent dopamine indicator dLight1.2 (AAV5-hSyn-dLight1.2) (Addgene, titer ≥ 4×10¹² vg/mL) and calcium indicator jRCaMP1b (AAV1.Syn.NES-jRCaMP1b.WPRE.SV40) (Addgene, titer ≥ 1×10¹³ vg/mL) were mixed in a 1:1 ratio and vortexed immediately prior to intracranial infusion surgeries.

### Drugs

Cocaine hydrochloride (graciously provided by NIDA) was dissolved at 2 mg/ml in 0.9% sterile saline (Thermo Fisher) and filter-sterilized.

### Behavioral Apparatus

Experiments were conducted in 4 standard operant conditioning chambers using MedPC software (Med Associates). Each animal underwent all training and testing in the same chamber. Each chamber was equipped with bar floors and a syringe pump connected to a swiveled leash. All chambers had 3 plexiglass walls and one wall containing two levers with cue lights above them, a head-entry magazine, a houselight, and a tone generator. Chambers were housed in a sound-attenuating box with a fan for background noise.

### Surgery

#### Anesthesia

Rats were fully anesthetized with ketamine (100 mg/kg, Henry Schein) and xylazine (5 mg/kg, Butler Schein) intramuscularly, administered analgesic (5 mg/kg Rimadyl, Henry Schein) subcutaneously, and prepared for surgery as previously described (*32*, *38*).

#### Viral infusion

Viral infusion surgery took place at least 4 weeks prior to photometry recordings to allow for virus expression. Rats were placed in a stereotaxic frame and lidocaine (0.3 ml, Butler Schein) was injected subcutaneously above the skull as previously described (*32*). A 26-gauge injection cannula connected to a Hamilton Syringe and pump was used to inject 1 μl of virus mixture at a rate of 0.05 μl/min unilaterally into the anterior DLS (in mm from bregma, anterior and posterior (AP): +0.8; medial and lateral (ML): ± 2.8; dorsal and ventral (DV): -5.0) and, in the opposite hemisphere, the posterior DMS (AP: -0.2 mm; ML: ± 2.2 mm; DV: -4.7 mm). The hemisphere receiving each injection was counterbalanced.

#### Intravenous catheterization and optic fiber implantation

In a second surgery up to a week before rats began self-administration, rats were implanted with a chronic indwelling intravenous catheter into the right jugular vein as previously described (*32*, *38*, *80*). Rats were then placed in a stereotaxic frame and lidocaine was injected subcutaneously above the skull. Fiber optic cannulae (Thorlabs, 2.5 mm ferrule, 400 μm core, 5 mm long) were lowered into the DLS (AP: +0.8 mm; ML: ± 3.0 mm; DV: -4.5 mm) and the DMS (AP: -0.2 mm; ML: ± 2.0 mm; DV: -4.0 mm) and were secured as previously described (*32*).

#### Post-operative care

Rats were administered analgesic and catheters were flushed daily to maintain patency as previously described (*38*).

### Behavioral Procedures

#### Cocaine self-administration

Rats were trained to self-administer cocaine (1mg/kg/infusion) in 1-hour daily sessions for 20 days as previously described (*38*). Briefly, cocaine infusions were paired with a 20-second audiovisual cue accompanied by a 20-second time-out when levers were retracted and the houselight was extinguished, and inactive lever presses were recorded but had no consequences. Rats were initially trained to self-administer cocaine on a fixed-ratio 1 (FR1) schedule for 7 days (acquisition) then on an FR3 schedule for 3 days (early training). Rats were then split into FR-trained (goal-directed) and SO-trained (habit-like) groups. FR-trained rats were maintained on an FR3 schedule for 5 days (middle training) and then trained on an FR5 schedule for 5 days (late training). SO-trained rats were trained for 5 days on an FR5(FR2S) schedule (middle training) followed by 5 days on an FR7(FR2S) schedule (late training). Details for SO schedule training were as previously described (*38*); briefly, every second lever press resulted in a 1-second presentation of the audiovisual cue (FR2S), and every fifth (FR5) or seventh (FR7) completion of the FR2S cycle resulted in cocaine infusion and timeout paired with the 20-second audiovisual cue. Levers were not retracted during 1-second cues during SO training.

#### Daily cue-induced drug-seeking tests

During fiber photometry recordings, rats underwent a 15-minute drug-seeking test immediately before self-administration on days 9-20. To prevent cocaine from influencing extracellular dopamine, no cocaine was administered during these tests, but audiovisual cues and timeouts occurred on the same reinforcement schedule each rat self-administered under on the previous day. Standard self-administration sessions immediately followed each of these 15-minute test sessions.

#### Pavlovian cue extinction

On the day immediately following the final day of self-administration, with levers retracted, rats were non-contingently exposed to 120 20-second audiovisual cues separated by 10 seconds during photometry recordings.

#### Cue-induced drug-seeking test

On the day following cue extinction, rats underwent a 1-hour drug-seeking test during fiber photometry recordings. Cocaine was withheld and cues and timeouts occurred on the rat’s previous reinforcement schedule. The cue-induced drug-seeking test was 1 hour long because, in our hands, rats can take longer to resume cue-induced drug seeking after cue extinction, which was supported by our finding that that multiple rats (n=4) did not complete their reinforcement schedule within the first 15 minutes of this drug-seeking test.

### Fiber Photometry

#### Recordings

Photometry recordings were collected using a multi-wavelength photometry system (Plexon) and a branched low-autofluorescence fiber-optic patch cord (Doric: 2 branches, 400 μm core, 440 μm cladding, 0.37 NA). Laser output for each excitation wavelength (560 nm, 465 nm, and 410 nm) was set that laser intensity at the cable tip was 20-30 μW. Laser was passed through the patch cord for 30 minutes prior to daily photometry recordings to minimize autofluorescence of the cable during recordings. Recordings occurred using 3-phase cycling of 415, 465, and 560 nm LEDs. Fluorescence data were collected at 30 frames per second using Plexon software, and behavioral events were aligned to photometry fluorescence using TTL timestamp outputs from MedPC software. Rats were habituated to the optic cable setup in the operant chamber for at least 1 day prior to photometry recordings.

#### Processing and analysis

Data were processed and analyzed using custom MATLAB (Mathworks, Natick, MA, USA) scripts, which are available upon request. Fluorescent signals were forward and reverse low-pass filtered at 30 Hz. Traces were visualized and motion artifacts were removed. The isosbestic control trace was fitted to fluorescent signal traces at 465 nm (dLight) and 560 nm (RCaMP) using a least squares polynomial fit of degree 1. ΔF/F was calculated by subtracting the fitted isosbestic signal from the fluorescent signal and then dividing by the fitted isosbestic signal. Z-scores were calculated across the entire fluorescent signal trace by subtracting the mean ΔF/F value from each data point and dividing by the standard deviation. Traces around behavioral events (including 3 seconds before and 30 seconds after) were separated and averaged for each animal for each recording day, and SEM was calculated for the average trace.

Because the majority of the responses we observed occurred within 1 second of behavioral events, the area under the curve (AUC) of the z-score and peak z-score amplitude in the 1 second post-event were calculated for all events. For each phase of training, each animal’s daily average z-score AUC and peak amplitude were averaged to obtain a single value per animal. For results, “cue-reinforced active lever presses” refer to lever presses that resulted in cue presentation and timeout (20-second audiovisual cue, houselight off, and lever retraction). “Unreinforced active lever presses” refer to active lever presses that did not result in any cue presentation, and for which there were no active lever presses (and therefore no cue presentations) in the 3 seconds before or 5 seconds after.

### Vaginal Cytology

Estrous cycle phase was determined daily by cellular morphology as previously described (*81*, *82*).

### Histology

Rats were anesthetized and perfused and brains were removed, sliced, mounted, and coverslipped as previously described (*32*). Every fourth section containing the dorsal striatum was mounted and imaged at 10X magnification using an Olympus BX61VS epifluorescent slide-scanning microscope to verify fiber placement location and virus expression at the base of the fiber.

### Exclusion Criteria

Rats were excluded from all analysis due to death or illness after surgery (n=5), loss of catheter patency (n=1) (determined by a 0.1 ml intravenous infusion of 10 mg/ml sodium brevital, Covetrus), or bilateral histological misses (n=4). For rats with a unilateral histological miss (n=6), data for the region with the histological miss were excluded. One rat was excluded from the final cue-induced drug-seeking test due to loss of head cap. Rats were excluded from analysis during cue-induced drug-seeking tests if they failed to make enough lever presses to reach a long cue during all phases of training (n=2). Rats were excluded from comparing activity during the late training to activity during drug seeking after cue extinction if they failed to obtain a long cue or make an isolated unreinforced lever press during either test (n=3).

### Quantification and Statistical Analysis

Behavioral data were collected using MedPC software. All statistical analyses were performed using GraphPad Prism. For the 1-hour cue-induced drug-seeking test after cue extinction, the ratio of responding was calculated by dividing the number of active lever presses during test by the number of active lever presses in the final self-administration session to normalize changes in responding across rats with different magnitudes of lever pressing behavior due to their training schedule.

For all statistical analyses, significance was set at p<0.05. All data were determined to be normally distributed using the Shapiro-Wilk test, and Bartlett’s test was used to determine that there were no significant differences in the estimated variance between groups. For analyses using repeated-measures ANOVAs, a Geisser-Greenhouse correction was used to account for potential lack of sphericity. Criteria for outlier data points was set at >2 standard deviations from the mean, and outlier points were excluded along with their paired data.

Infusions were analyzed by two-way rmANOVA, using time and training schedule as factors. Lever presses during self-administration were analyzed by three-way rmANOVA with time, training schedule, and lever as factors. Ratio of active lever presses during the post-cue extinction cue-induced drug-seeking test was analyzed with an unpaired student’s t-test. Calcium and dopamine peak z-score amplitude and AUCs during cue-induced drug-seeking tests were analyzed by two-way or three-way ANOVA with future training schedule, training schedule, sex, cue reinforcement, cue length, phase of training, or cue extinction as factors as indicated. Calcium and dopamine peak z-score amplitude and AUC during early training were analyzed to determine if there was an effect of estrous phase on signals by mixed-effects analysis with estrous phase and cue reinforcement as factors because one rat was never in estrus during those three days of testing. Calcium and dopamine peak z-score amplitude and AUCs during cue extinction were analyzed by one-way rmANOVA. For correlation analyses, Pearson’s correlation coefficients were calculated with average active lever presses as the independent variable and peak z-score amplitude or AUC as the dependent variable. Calcium and dopamine peak z-score amplitude and AUCs after pre-learning stimuli exposure were analyzed by two-way ANOVA with stimulus type and stimulus presentation as factors. When a significant effect was detected by one-way ANOVA or an interaction was detected by two-way or three-way ANOVA analysis, significant effects were further analyzed by Tukey’s or Sidak’s post-hoc multiple comparisons analysis, respectively. Throughout our analyses, we report the peak z-score amplitude as our primary measure of calcium and dopamine responses. Results from analyses of AUC data were overall similar and are therefore not presented in the main text, but there were some minor differences, so AUC results are presented in the supplementary material.

## Acknowledgments

We would like to acknowledge Dana Smith, Camryn Forbes, Michael Wright, Alexis Egazarian, Lauren Charlton, Lindsey Buchman, Alina Owsiany, and Juan Robayo for assistance with behavioral experiments, surgical procedures, and histology; Susanne Ahmari for technical advice; and Zoe LaPalombara for providing the initial code we adapted for data analysis.

## Author contributions

Conceptualization: BNB, MMT

Methodology: BNB, SJS, MMT

Investigation: BNB

Visualization and Analysis: BNB, SJS

Supervision: MMT

Writing—original draft: BNB

Writing—review & editing: BNB, SJS, MMT

## Data and materials availability

All data, code, and materials used in the analyses will be made available upon request.

## References

1. B. L. Carter, S. T. Tiffany, Meta-analysis of cue-reactivity in addiction research. Addiction. 94, 327–40 (1999).

2. S. Grant, E. D. London, D. B. Newlin, V. L. Villemagne, X. Liu, C. Contoreggi, R. L. Phillips, A. S. Kimes, A. Margolin, Activation of memory circuits during cue-elicited cocaine craving. Proc. Natl. Acad. Sci. 93, 12040–12045 (1996).

3. A. L. Milton, B. J. Everitt, The persistence of maladaptive memory: Addiction, drug memories and anti-relapse treatments. Neurosci. Biobehav. Rev. 36, 1119–1139 (2012).

4. G. J. Wang, N. D. Volkow, J. S. Fowler, P. Cervany, R. J. Hitzemann, N. R. Pappas, C. T. Wong, C. Felder, Regional brain metabolic activation during craving elicited by recall of previous drug experiences. Life Sci. 64, 775–84 (1999).

5. K. H. MacNiven, E. L. S. Jensen, N. Borg, C. B. Padula, K. Humphreys, B. Knutson, Association of Neural Responses to Drug Cues With Subsequent Relapse to Stimulant Use. JAMA Netw. open. 1, e186466 (2018).

6. M. M. Torregrossa, P. R. Corlett, J. R. Taylor, Aberrant learning and memory in addiction. Neurobiol. Learn. Mem. 96, 609–623 (2011).

7. T. M. Furlong, L. H. Corbit, R. A. Brown, B. W. Balleine, Methamphetamine promotes habitual action and alters the density of striatal glutamate receptor and vesicular proteins in dorsal striatum. Addict. Biol. 23, 857–867 (2018).

8. A. Nelson, S. Killcross, Amphetamine exposure enhances habit formation. J. Neurosci. 26, 3805–12 (2006).

9. R. E. Nordquist, P. Voorn, J. G. de Mooij-van Malsen, R. N. J. M. A. Joosten, C. M. A. Pennartz, L. J. M. J. Vanderschuren, Augmented reinforcer value and accelerated habit formation after repeated amphetamine treatment. Eur. Neuropsychopharmacol. 17, 532–40 (2007).

10. B. J. Everitt, T. W. Robbins, Neural systems of reinforcement for drug addiction: from actions to habits to compulsion. Nat. Neurosci. 8, 1481–1489 (2005).

11. B. N. Bender, M. M. Torregrossa, Molecular and circuit mechanisms regulating cocaine memory. Cell. Mol. Life Sci. 77, 3745–3768 (2020).

12. P. Olausson, J. D. Jentsch, D. D. Krueger, N. C. Tronson, A. C. Nairn, J. R. Taylor, Orbitofrontal cortex and cognitive-motivational impairments in psychostimulant addiction: evidence from experiments in the non-human primate. Ann. N. Y. Acad. Sci. 1121, 610–638 (2007).

13. B. J. Everitt, T. W. Robbins, From the ventral to the dorsal striatum: devolving views of their roles in drug addiction. Neurosci. Biobehav. Rev. 37, 1946–54 (2013).

14. S. B. Ostlund, B. W. Balleine, On habits and addiction: an associative analysis of compulsive drug seeking (NIH Public Access, 2008; http://www.ncbi.nlm.nih.gov/pubmed/20582332), vol. 5.

15. R. J. Smith, L. S. Laiks, Behavioral and neural mechanisms underlying habitual and compulsive drug seeking. Prog. Neuro-Psychopharmacology Biol. Psychiatry. 87, 11–21 (2017).

16. A. J. Gruber, R. J. McDonald, Context, emotion, and the strategic pursuit of goals: interactions among multiple brain systems controlling motivated behavior. Front. Behav. Neurosci. 6, 50 (2012).

17. K.-C. Leong, C. R. Berini, S. M. Ghee, C. M. Reichel, Extended cocaine-seeking produces a shift from goal-directed to habitual responding in rats. Physiol. Behav. 164, 330–5 (2016).

18. M. W. Shiflett, B. W. Balleine, Molecular substrates of action control in cortico-striatal circuits. Prog. Neurobiol. 95, 1–13 (2011).

19. N. D. Volkow, G. J. Wang, F. Telang, J. S. Fowler, J. Logan, A. R. Childress, M. Jayne, Y. Ma, C. Wong, Cocaine cues and dopamine in dorsal striatum: Mechanism of craving in cocaine addiction. J. Neurosci. 26, 6583–6588 (2006).

20. H. Garavan, J. Pankiewicz, A. Bloom, J. K. Cho, L. Sperry, T. J. Ross, B. J. Salmeron, R. Risinger, D. Kelley, E. A. Stein, Cue-induced cocaine craving: neuroanatomical specificity for drug users and drug stimuli. Am. J. Psychiatry. 157, 1789–1798 (2000).

21. L. C. Maas, S. E. Lukas, M. J. Kaufman, R. D. Weiss, S. L. Daniels, V. W. Rogers, T. J. Kukes, P. F. Renshaw, Functional magnetic resonance imaging of human brain activation during cue-induced cocaine craving. Am. J. Psychiatry. 155, 124–126 (1998).

22. B. E. Wexler, C. H. Gottschalk, R. K. Fulbright, I. Prohovnik, C. M. Lacadie, B. J. Rounsaville, J. C. Gore, Functional magnetic resonance imaging of cocaine craving. Am. J. Psychiatry. 158, 86–95 (2001).

23. T. R. Kosten, B. E. Scanley, K. A. Tucker, A. Oliveto, C. Prince, R. Sinha, M. N. Potenza, P. Skudlarski, B. E. Wexler, Cue-induced brain activity changes and relapse in cocaine-dependent patients. Neuropsychopharmacology. 31, 644–650 (2006).

24. J. J. Prisciandaro, H. Myrick, S. Henderson, A. L. McRae-Clark, K. T. Brady, Prospective associations between brain activation to cocaine and no-go cues and cocaine relapse. Drug Alcohol Depend. 131, 44–49 (2013).

25. A. Milton, Drink, drugs and disruption: memory manipulation for the treatment of addiction. Curr. Opin. Neurobiol. 23, 706–712 (2013).

26. C. A. Conklin, S. T. Tiffany, Applying extinction research and theory to cue-exposure addiction treatments. Addiction. 97, 155–67 (2002).

27. M. M. Torregrossa, J. R. Taylor, Learning to forget: manipulating extinction and reconsolidation processes to treat addiction. Psychopharmacology (Berl). 226, 659–72 (2013).

28. M. B. Powers, P. M. G. Emmelkamp, Virtual reality exposure therapy for anxiety disorders: A meta-analysis. J. Anxiety Disord. 22, 561–569 (2008).

29. M. B. Powers, J. M. Halpern, M. P. Ferenschak, S. J. Gillihan, E. B. Foa, A meta-analytic review of prolonged exposure for posttraumatic stress disorder. Clin. Psychol. Rev. 30, 635–641 (2010).

30. H. B. Madsen, I. C. Zbukvic, S. J. Luikinga, A. J. Lawrence, J. H. Kim, Extinction of conditioned cues attenuates incubation of cocaine craving in adolescent and adult rats. Neurobiol. Learn. Mem. 143, 88–93 (2017).

31. C. J. Perry, F. Reed, I. C. Zbukvic, J. H. Kim, A. J. Lawrence, The metabotropic glutamate 5 receptor is necessary for extinction of cocaine-associated cues. Br. J. Pharmacol. 173, 1085–1094 (2016).

32. M. T. Rich, Y. H. Huang, M. M. Torregrossa, Plasticity at Thalamo-amygdala Synapses Regulates Cocaine-Cue Memory Formation and Extinction. Cell Rep. 26, 1010–1020.e5 (2019).

33. M. M. Torregrossa, J. Gordon, J. R. Taylor, Double Dissociation between the Anterior Cingulate Cortex and Nucleus Accumbens Core in Encoding the Context versus the Content of Pavlovian Cocaine Cue Extinction. J. Neurosci. 33, 8370–8377 (2013).

34. A. I. Mellentin, L. Skøt, B. Nielsen, G. M. Schippers, A. S. Nielsen, E. Stenager, C. Juhl, Cue exposure therapy for the treatment of alcohol use disorders: A meta-analytic review. Clin. Psychol. Rev. 57, 195–207 (2017).

35. M. M. Torregrossa, H. Sanchez, J. R. Taylor, D-cycloserine reduces the context specificity of pavlovian extinction of cocaine cues through actions in the nucleus accumbens. J. Neurosci. 30, 10526–33 (2010).

36. M. T. Rich, M. M. Torregrossa, Molecular and synaptic mechanisms regulating drug-associated memories: Towards a bidirectional treatment strategy. Brain Res. Bull. 141, 58–71 (2018).

37. K. M. Kantak, B. Á. Nic Dhonnchadha, Pharmacological enhancement of drug cue extinction learning: translational challenges. Ann. N. Y. Acad. Sci. 1216, 122–37 (2011).

38. B. N. Bender, M. M. Torregrossa, Dorsolateral striatum dopamine-dependent cocaine seeking is resistant to pavlovian cue extinction in male and female rats. Neuropharmacology. 182 (2021), doi:10.1016/J.NEUROPHARM.2020.108403.

39. L. H. Corbit, H. Nie, P. H. Janak, Habitual alcohol seeking: time course and the contribution of subregions of the dorsal striatum. Biol. Psychiatry. 72, 389–95 (2012).

40. B. J. Knowlton, T. K. Patterson, Habit formation and the striatum. Curr. Top. Behav. Neurosci. 37, 275–295 (2018).

41. J. M. Barker, L. H. Corbit, D. L. Robinson, C. M. Gremel, R. A. Gonzales, L. J. Chandler, Corticostriatal circuitry and habitual ethanol seeking. Alcohol. 49, 817–824 (2015).

42. J. E. Murray, D. Belin, B. J. Everitt, Double dissociation of the dorsomedial and dorsolateral striatal control over the acquisition and performance of cocaine seeking. Neuropsychopharmacology. 37, 2456–66 (2012).

43. D. Belin, B. J. Everitt, Cocaine seeking habits depend upon dopamine-dependent serial connectivity linking the ventral with the dorsal striatum. Neuron. 57, 432–41 (2008).

44. A. Faure, U. Haberland, F. Condé, N. El Massioui, Lesion to the Nigrostriatal Dopamine System Disrupts Stimulus-Response Habit Formation. J. Neurosci. 25, 2771–2780 (2005).

45. C. M. Gremel, R. M. Costa, Orbitofrontal and striatal circuits dynamically encode the shift between goal-directed and habitual actions. Nat. Commun. 4, 2264 (2013).

46. S. Kato, R. Fukabori, K. Nishizawa, K. Okada, N. Yoshioka, M. Sugawara, Y. Maejima, K. Shimomura, M. Okamoto, S. Eifuku, K. Kobayashi, Action Selection and Flexible Switching Controlled by the Intralaminar Thalamic Neurons. Cell Rep. 22, 2370–2382 (2018).

47. K. K. Cover, U. Gyawali, W. G. Kerkhoff, M. H. Patton, C. Mu, M. G. White, A. E. Marquardt, B. M. Roberts, J. F. Cheer, B. N. Mathur, Activation of the Rostral Intralaminar Thalamus Drives Reinforcement through Striatal Dopamine Release. Cell Rep. 26, 1389–1398.e3 (2019).

48. S. Killcross, E. Coutureau, Coordination of actions and habits in the medial prefrontal cortex of rats. Cereb. Cortex. 13, 400–8 (2003).

49. N. W. Lingawi, B. W. Balleine, Amygdala central nucleus interacts with dorsolateral striatum to regulate the acquisition of habits. J. Neurosci. 32, 1073–1081 (2012).

50. J. E. Murray, A. Belin-Rauscent, M. Simon, C. Giuliano, M. Benoit-Marand, B. J. Everitt, D. Belin, Basolateral and central amygdala differentially recruit and maintain dorsolateral striatum-dependent cocaine-seeking habits. Nat. Commun. 6, 10088 (2015).

51. T. A. Shnitko, D. L. Robinson, Regional variation in phasic dopamine release during alcohol and sucrose self-administration in rats. ACS Chem. Neurosci. 6, 147–154 (2015).

52. R. Ito, J. W. Dalley, T. W. Robbins, B. J. Everitt, Dopamine release in the dorsal striatum during cocaine-seeking behavior under the control of a drug-associated cue. J. Neurosci. 22, 6247–53 (2002).

53. M. Klanker, L. Fellinger, M. Feenstra, I. Willuhn, D. Denys, Regionally distinct phasic dopamine release patterns in the striatum during reversal learning. Neuroscience. 345, 110–123 (2017).

54. I. Willuhn, L. M. Burgeno, B. J. Everitt, P. E. M. Phillips, Hierarchical recruitment of phasic dopamine signaling in the striatum during the progression of cocaine use. Proc. Natl. Acad. Sci. U. S. A. 109, 20703–20708 (2012).

55. H. D. Brown, J. E. Mccutcheon, J. J. Cone, M. E. Ragozzino, M. F. Roitman, Primary food reward and reward predictive stimuli evoke different patterns of phasic dopamine signaling throughout the striatum. Eur. J. Neurosci. 34, 1997 (2011).

56. R. R. Fanelli, J. T. Klein, R. M. Reese, D. L. Robinson, Dorsomedial and dorsolateral striatum exhibit distinct phasic neuronal activity during alcohol self-administration in rats. Eur. J. Neurosci. 38, 2637–2648 (2013).

57. E. Y. Kimchi, M. M. Torregrossa, J. R. Taylor, M. Laubach, Neuronal Correlates of Instrumental Learning in the Dorsal Striatum. J. Neurophysiol. 102, 475 (2009).

58. Y. Vandaele, N. R. Mahajan, D. J. Ottenheimer, J. M. Richard, S. P. Mysore, P. H. Janak, Distinct recruitment of dorsomedial and dorsolateral striatum erodes with extended training. Elife. 8 (2019), doi:10.7554/eLife.49536.

59. Y. Vandaele, P. H. Janak, Lack of action monitoring as a prerequisite for habitual and chunked behavior: Behavioral and neural correlates. iScience. 26 (2022), doi:10.1016/J.ISCI.2022.105818.

60. T. Patriarchi, J. R. Cho, K. Merten, M. W. Howe, A. Marley, W. H. Xiong, R. W. Folk, G. J. Broussard, R. Liang, M. J. Jang, H. Zhong, D. Dombeck, M. von Zastrow, A. Nimmerjahn, V. Gradinaru, J. T. Williams, L. Tian, Ultrafast neuronal imaging of dopamine dynamics with designed genetically encoded sensors. Science (80-.). 360 (2018), doi:10.1126/science.aat4422.

61. T. Patriarchi, J. R. Cho, K. Merten, A. Marley, G. J. Broussard, R. Liang, J. Williams, A. Nimmerjahn, M. von Zastrow, V. Gradinaru, L. Tian, Imaging neuromodulators with high spatiotemporal resolution using genetically encoded indicators. Nat. Protoc. 14, 3471–3505 (2019).

62. Y. Wang, E. M. DeMarco, L. S. Witzel, J. D. Keighron, A selected review of recent advances in the study of neuronal circuits using fiber photometry. Pharmacol. Biochem. Behav. 201, 173113 (2021).

63. Y. Li, Z. Liu, Q. Guo, M. Luo, Long-term Fiber Photometry for Neuroscience Studies. Neurosci. Bull. 35, 425–433 (2019).

64. C. A. Siciliano, K. M. Tye, Leveraging calcium imaging to illuminate circuit dysfunction in addiction. Alcohol. 74 (2019), pp. 47–63.

65. A. A. Legaria, B. A. Matikainen-Ankney, B. Yang, B. Ahanonu, J. A. Licholai, J. G. Parker, A. V. Kravitz, Fiber photometry in striatum reflects primarily nonsomatic changes in calcium. Nat. Neurosci. 2022, 1–5 (2022).

66. J. Mejaes, D. Desai, C. A. Siciliano, D. J. Barker, Practical opinions for new fiber photometry users to obtain rigorous recordings and avoid pitfalls. Pharmacol. Biochem. Behav. 221, 173488 (2022).

67. L. H. Corbit, B. C. Chieng, B. W. Balleine, Effects of Repeated Cocaine Exposure on Habit Learning and Reversal by N-Acetylcysteine. Neuropsychopharmacology. 39, 1893–1901 (2014).

68. H. H. Yin, S. B. Ostlund, B. J. Knowlton, B. W. Balleine, The role of the dorsomedial striatum in instrumental conditioning. Eur. J. Neurosci. 22, 513–523 (2005).

69. P. Watson, S. de Wit, Current limits of experimental research into habits and future directions. Curr. Opin. Behav. Sci. 20, 33–39 (2018).

70. J. Morrens, Ç. Aydin, A. Janse van Rensburg, J. Esquivelzeta Rabell, S. Haesler, Cue-Evoked Dopamine Promotes Conditioned Responding during Learning. Neuron. 106, 142–153.e7 (2020).

71. S. B. Flagel, J. J. Clark, T. E. Robinson, L. Mayo, A. Czuj, I. Willuhn, C. A. Akers, S. M. Clinton, P. E. M. Phillips, H. Akil, A selective role for dopamine in reward learning. Nature. 469, 53 (2011).

72. J. L. Seiler, C. V. Cosme, V. N. Sherathiya, M. D. Schaid, J. M. Bianco, A. S. Bridgemohan, T. N. Lerner, Dopamine signaling in the dorsomedial striatum promotes compulsive behavior. Curr. Biol. 32, 1175–1188.e5 (2022).

73. D. A. Kupferschmidt, K. Juczewski, G. Cui, K. A. Johnson, D. M. Lovinger, Parallel, but Dissociable, Processing in Discrete Corticostriatal Inputs Encodes Skill Learning. Neuron. 96, 476–489.e5 (2017).

74. K. S. Zimmermann, J. A. Yamin, D. G. Rainnie, K. J. Ressler, S. L. Gourley, Connections of the Mouse Orbitofrontal Cortex and Regulation of Goal-Directed Action Selection by Brain-Derived Neurotrophic Factor. Biol. Psychiatry. 81, 366–377 (2017).

75. K. D. Alloway, J. B. Smith, T. M. Mowery, G. D. R. Watson, Sensory Processing in the Dorsolateral Striatum: The Contribution of Thalamostriatal Pathways. Front. Syst. Neurosci. 11, 53 (2017).

76. E. N. Holly, M. F. Davatolhagh, K. Choi, O. O. Alabi, L. Vargas Cifuentes, M. V. Fuccillo, Striatal Low-Threshold Spiking Interneurons Regulate Goal-Directed Learning. Neuron. 103, 92–101.e6 (2019).

77. J. K. O’Hare, H. Li, N. Kim, E. Gaidis, K. Ade, J. Beck, H. Yin, N. Calakos, Striatal fast-spiking interneurons selectively modulate circuit output and are required for habitual behavior. Elife. 6 (2017), doi:10.7554/eLife.26231.

78. E. Garr, A. R. Delamater, Chemogenetic inhibition in the dorsal striatum reveals regional specificity of direct and indirect pathway control of action sequencing. Neurobiol. Learn. Mem. 169, 107169 (2020).

79. H. H. Yin, S. P. Mulcare, M. R. F. Hilário, E. Clouse, T. Holloway, M. I. Davis, A. C. Hansson, D. M. Lovinger, R. M. Costa, Dynamic reorganization of striatal circuits during the acquisition and consolidation of a skill. Nat. Neurosci. 12, 333–341 (2009).

80. M. M. Torregrossa, P. W. Kalivas, Neurotensin in the ventral pallidum increases extracellular gamma-aminobutyric acid and differentially affects cue- and cocaine-primed reinstatement. J. Pharmacol. Exp. Ther. 325, 556–66 (2008).

81. J. N. Parrish, M. L. Bertholomey, H. W. Pang, R. C. Speth, M. M. Torregrossa, Estradiol modulation of the renin– angiotensin system and the regulation of fear extinction. Transl. Psychiatry. 9 (2019), doi:10.1038/S41398-019-0374-0.

82. B. N. Bender, M. M. Torregrossa, Intermittent cocaine self-administration has sex-specific effects on addiction-like behaviors in rats. Neuropharmacology. 230, 109490 (2023).

