## supplement for "Changes in dorsomedial striatum activity mediate expression of goal-directed vs. habit-like cue-induced cocaine seeking"

**Supplementary Text**

**There is no effect of sex or estrous phase on dorsal striatal calcium or dopamine responses to lever presses**

Because we used both male and female rats in these experiments, we wanted to determine if there were any effects of sex or estrous phase on dorsal striatal activity during drug seeking. During the early phase of training, before rats were split into FR- and SO-trained groups, we compared activity between males and females using 2-way ANOVAs. For DLS calcium peak amplitude, there was a main effect of cue reinforcement ($F_{(1,10)}$=10.96, p=0.0079), but no effect of sex ($F_{(1,10)}$=3.416, p=0.0943) or cue reinforcement × sex interaction ($F_{(1,10)}$=2.403, p=0.1521) (Figure S1A). Similarly, for DMS calcium peak amplitude, there was a main effect of cue reinforcement ($F_{(1,11)}$=12.42, p=0.0048), but no effect of sex ($F_{(1,11)}$=3.703, p=0.0806) or cue reinforcement × sex interaction ($F_{(1,11)}$=0.7044, p=0.4192) (Figure S1B). There was no main effect of cue reinforcement ($F_{(1,11)}$=0.003257, p=0.9555; $F_{(1,11)}$=0.0007764, p=0.9103) or sex ($F_{(1,11)}$=0.4337, p=0.5237; $F_{(1,11)}$=0.02260, p=0.6880) or cue reinforcement × sex interaction ($F_{(1,11)}$=0.5442, p=0.4761; $F_{(1,11)}$=0.1285, p=0.7268) for DLS (Figure S1C) or DMS (Figure S1D) dopamine peak amplitude. There was a main effect of cue reinforcement ($F_{(1,10)}$=36.38, p=0.0001; $F_{(1,11)}$=23.37, p=0.0005), but no effect of sex ($F_{(1,10)}$=0.9574, p=0.3509; $F_{(1,11)}$=1.860, p=0.1998) or cue reinforcement × sex interaction ($F_{(1,10)}$=0.3662, p=0.5585; $F_{(1,11)}$=1.865, p=0.1993) for DLS (Figure S1E) and DMS (Figure S1F) calcium AUC. There was no main effect of cue reinforcement ($F_{(1,11)}$=4.356, p=0.0609; $F_{(1,11)}$=2.843, p=0.1199) or sex ($F_{(1,11)}$=0.3945, p=0.5428; $F_{(1,11)}$=0.02097, p=0.8875) or cue reinforcement × sex interaction ($F_{(1,11)}$=0.3112, p=0.1277; $F_{(1,11)}$= 0.3112, p=0.5881) for DLS (Figure S1G) or DMS (Figure S1H) dopamine AUC.

Most females (n=6 of 7) were in estrus during one of the 3 days of the early training phase, so we also examined if calcium or dopamine responses to lever presses differed when females were in estrus or another phase of the estrous cycle (non-estrus). For these analyses, mixed effects analyses were used to accommodate one rat not being in estrus and another not making a cue-reinforced lever press while in estrus. For DLS (Figure S1I) and DMS (Figure S1J) calcium peak amplitude, there was a main effect of cue reinforcement ($F_{(1,5)}$=9.646, p=0.0267; $F_{(1,6)}$=8.109, p=0.0293), but no effect of estrous phase ($F_{(1,5)}$=0.01962, p=0.8941; $F_{(1,6)}$=0.5886, p=0.4721) or cue reinforcement × estrous phase interaction ($F_{(1,2)}$=2.386, p=0.2624; $F_{(1,3)}$=2.399, p=0.2192). For DLS (Figure S1K) and DMS (Figure S1L) dopamine peak amplitude, there was no main effect of cue reinforcement ($F_{(1,5)}$=0.5567, p=0.4892; $F_{(1,6)}$=0.0004565, p=0.9836) or estrous phase ($F_{(1,5)}$=0.1196, p=0.7436; $F_{(1,6)}$=2.614, p=0.1570) or cue reinforcement × estrous phase interaction ($F_{(1,2)}$=11.41, p=0.0776; $F_{(1,3)}$=0.3294, p=0.6062). For DLS (Figure S1M) and DMS (Figure S1N) calcium AUC, there was a main effect of cue reinforcement ($F_{(1,5)}$=26.75, p=0.0035; $F_{(1,6)}$=20.64, p=0.0039), but no effect of estrous phase ($F_{(1,5)}$=0.1281, p=0.7351; $F_{(1,6)}$=0.4512, p=0.5268) or cue reinforcement × estrous phase interaction ($F_{(1,2)}$=1.252, p=0.3795; $F_{(1,3)}$=1.870, p=0.2649). There was no main effect of cue reinforcement ($F_{(1,5)}$=0.3965, p=0.5565; $F_{(1,21)}$=3.932, p=0.0606) or estrous phase ($F_{(1,5)}$=0.3025, p=0.6060; $F_{(1,21)}$=0.05453, p=0.8176) or cue reinforcement × estrous phase interaction ($F_{(1,2)}$=3.826, p=0.1896; $F_{(1,21)}$=0.2593, p=0.6159) for DLS (Figure S1O) and DMS (Figure S1P) dopamine AUC. Overall, these data indicate that there was no effect of sex or estrous phase on dorsal striatal calcium or dopamine responses to lever presses, and we therefore collapsed results across sex throughout analyses in the main text.

**During early training, cue-reinforced lever presses result in increased AUC for DLS calcium and dopamine and DMS calcium**

During early training, prior to being split into FR- and SO-trained groups, we found that calcium peak amplitude in both the DLS and DMS was greater for cue-reinforced lever presses, but this was not the case for dopamine peak amplitude (Figure 2). Additionally, there were no differences in peak amplitude between rats that would later be split into FR- and SO-trained groups (Figure 2). In addition to peak amplitude data, we also calculated the AUC in the 1 second after lever press. Results were overall similar, with a few differences. There was a main effect of cue reinforcement ($F_{(1,8)}$=35.23, p=0.0003) as well as future training schedule ($F_{(1,8)}$=7.727, p=0.0239) for DLS calcium AUC, but no interaction ($F_{(1,8)}$=0.3938, p=0.5478) (2-way ANOVA) (Figure S2A). These data suggest that rats later placed in the SO-trained group may have by chance had greater calcium responses to lever presses in the DLS during early training (rats were split into groups randomly and photometry analysis occurred at the end of the experiment). Importantly, this difference was not present for peak analysis (Figure 2A) and did not persist throughout late training, and this difference does not greatly impact our overall interpretation of results. For DMS calcium AUC, there was a main effect of cue reinforcement ($F_{(1,9)}$=16.81, p=0.0027), but no effect of future training schedule ($F_{(1,9)}$=0.05188, p=0.8249) or interaction ($F_{(1,9)}$=0.2703, p=0.6157) (Figure S2B). For DLS dopamine AUC, there was a main effect of cue reinforcement ($F_{(1,9)}$=7.931, p=0.0202), but no effect of future training schedule ($F_{(1,9)}$=1.017, p=0.3397) or interaction ($F_{(1,9)}$=2.256, p=0.1673) (Figure S2C). The effect of cue was not present for DLS dopamine peak amplitude during early training (Figure 2C), likely because peak analysis isn't sensitive to the mainly negative AUC in response to unreinforced lever presses shown here. There was no main effect of cue ($F_{(1,9)}$=2.516, p=0.1513) or future training schedule ($F_{(1,9)}$=1.282, p=0.2903) or interaction ($F_{(1,9)}$=4.229, p=0.0738) for DMS dopamine AUC (Figure S2D). Though only statistically significant for the DLS, these results suggest that in this early training phase there appears to be a trend toward a reduction in dorsal striatal dopamine in response to unreinforced lever presses. Interestingly, these recordings took place during the first few days of FR3 training, when animals were for the first time making active lever presses that did not result in cue reinforcement, which previously occurred on an FR1 schedule. Therefore, these results may reflect an aspect of reward prediction error, where a lever press that didn't result in the expected reinforcing outcome resulted in a reduction in dorsal striatal dopamine.

**Average daily lever presses during the early phase of training are not correlated with calcium or dopamine responses to cue-reinforced lever presses**

Because rats were later trained on schedules of reinforcement that required increasing numbers of lever presses, we wanted to determine if there was a correlation between number of lever presses and dorsal striatal calcium and dopamine activity. We found no correlation between average daily active lever presses during early training and DLS calcium peak amplitude ($r=-0.1212$, $p=0.7213$) (Figure S2E), DMS calcium peak amplitude ($r=0.07310$, $p=0.8309$) (Figure S2F), DLS dopamine peak amplitude ($r=0.2325$, $p=0.4914$) (Figure S2G), DMS dopamine peak amplitude ($r=-0.5745$, $p=0.0645$) (Figure S2H), DLS calcium AUC ($r=-0.1935$, $p=0.5686$) (Figure S2I), DMS calcium AUC ($r=0.08215$, $p=0.8102$) (Figure S2J), DLS dopamine AUC ($r=0.3760$, $p=0.2544$) (Figure S2K), or DMS dopamine AUC ($r=-0.2970$, $p=0.3751$) (Figure S2L) (Pearson's correlation).

**Dorsal striatal calcium and dopamine AUC after cue-reinforced lever presses differ between FR-trained and SO-trained rats**

During middle and late training, AUC in the 1 second after lever presses for calcium and dopamine in the DLS and DMS were compared between groups across training phases using 3-way ANOVAs. For DLS calcium AUC, there was a main effect of cue reinforcement ($F_{(1,9)}=20.34$, $p=0.0015$) and training schedule ($F_{(1,9)}=6.695$, $p=0.0293$) as well as a training schedule × phase of training interaction ($F_{(1,9)}=6.972$, $p=0.0269$), but there was no main effect of phase of training ($F_{(1,9)}=3.892$, $p=0.0821$) or cue reinforcement × training schedule ($F_{(1,9)}=2.381$, $p=0.1572$), cue reinforcement × phase of training ($F_{(1,9)}=0.9052$, $p=0.3662$) or 3-way interaction ($F_{(1,9)}=2.882$, $p=0.1238$) (Figure S3A). These results slightly differ from those for DLS calcium peak amplitude, where there was only a main effect of cue reinforcement (Figure 3A), and suggest that SO-trained rats had a greater DLS calcium AUC for cue-reinforced and unreinforced lever presses than FR-trained rats during the middle phase of training. However, this difference cannot necessarily be attributed to SO training, because these rats also showed greater calcium AUC responses to lever presses during early training, before training schedules differed between groups. For DMS calcium AUC, there was a main effect of cue reinforcement ($F_{(1,9)}$=9.318, p=0.0137) as well as a training schedule × phase of training interaction ($F_{(1,9)}$=20.77, p=0.0014), but no main effects of training schedule ($F_{(1,9)}$=0.01153, p=0.9168), phase of training ($F_{(1,9)}$=2.825, p=0.1271) or cue reinforcement × training schedule ($F_{(1,9)}$=4.775, p=0.0567), cue reinforcement × phase of training ($F_{(1,9)}$=0.03256, p=0.8608) or 3-way interaction ($F_{(1,9)}$=0.05292, p=0.8076) (Figure S3B). These results suggest that although there was an overall effect of cue reinforcement on DMS calcium AUC for both groups, DMS calcium AUC after any lever press was reduced in SO-trained rats in the late phase of training. Although these results slightly differ than those for DMS calcium peak amplitude (Figure 3B), the overall interpretation that DMS calcium activity was reduced after extended SO training is supported.

For DLS dopamine AUC, there was a main effect of cue reinforcement ($F_{(1,9)}$=13.67, p=0.0049) as well as a cue reinforcement × phase of training ($F_{(1,9)}$=8.994, p=0.0150) and training schedule × phase of training interaction ($F_{(1,9)}$=6.873, p=0.0277), but no main effect of training schedule ($F_{(1,9)}$=2.696, p=0.1350) or phase of training ($F_{(1,9)}$=0.2766, p=0.6117), and there were no cue reinforcement × training schedule ($F_{(1,9)}$=1.648, p=0.2313) or 3-way interactions ($F_{(1,9)}$=0.2454, p=0.6322) (Figure S3C). These data suggest that while both groups showed enhanced DLS dopamine AUC to cue-reinforced lever presses versus unreinforced lever presses during the middle phase of training, overall DLS dopamine AUC was greater for SO-trained rats in the middle phase of training. DLS dopamine AUC results differed from those for DLS dopamine peak (Figure 3C), which suggest that DLS dopamine peak amplitude may be more related to SO-training, whereas DLS dopamine AUC may be more related to phase of training as well as training schedule. For DMS dopamine AUC, there was no main effect of cue reinforcement ($F_{(1,9)}$=3.556, p=0.0920), training schedule ($F_{(1,9)}$=0.1157, p=0.7416), or phase of training ($F_{(1,9)}$=0.06244, p=0.8083), and there were no cue reinforcement × training schedule ($F_{(1,9)}$=0.02924, p=0.8680), cue reinforcement × phase of training ($F_{(1,9)}$=0.1923, p=0.6713), training schedule × phase of training ($F_{(1,9)}$=0.1933, p=0.6705), or 3-way interactions ($F_{(1,9)}$=0.2752, p=0.6125) (Figure S3D).

**Cue extinction does not impact dorsal striatal calcium or dopamine AUC responses to lever presses**

Dorsal striatal calcium and dopamine AUC in response to cue-reinforced and unreinforced active lever presses were compared during drug-seeking tests during the late phase of training (pre-ext) and after cue extinction (post-ext) using 3-way ANOVAs. For DLS calcium AUC, there was a main effect of cue reinforcement ($F_{(1,7)}$=14.26, p=0.0069) and training schedule ($F_{(1,7)}$=11.47, p=0.0117), but there was no main effect of cue extinction ($F_{(1,7)}$=0.8767, p=0.3803) or cue reinforcement × training schedule ($F_{(1,7)}$=0.2842, p=0.6105), cue reinforcement × cue extinction ($F_{(1,7)}$=0.03958, p=0.8480), training schedule × cue extinction ($F_{(1,7)}$=3.445, p=0.1058), or 3-way interaction ($F_{(1,7)}$=0.2140, p=0.6577) (Figure S4A). For DMS calcium AUC, there was a main effect of cue reinforcement ($F_{(1,7)}$=42.11, p=0.0003) and there was also a cue reinforcement × training schedule interaction ($F_{(1,7)}$=24.83, p=0.0016), but there were no main effects of training schedule ($F_{(1,7)}$=1.783, p=0.2235) or cue extinction ($F_{(1,7)}$=1.545, p=0.2535) or cue reinforcement × cue extinction ($F_{(1,7)}$=0.1873, p=0.6782), training schedule × cue extinction ($F_{(1,7)}$=4.800, p=0.0646), or 3-way interactions ($F_{(1,7)}$=0.001162, p=0.9738) (Figure S4B). For both DLS (Figure S4C) and DMS (Figure S4D) dopamine AUC, there was a main effect of cue reinforcement ($F_{(1,7)}$=9.386, p=0.0182; $F_{(1,7)}$=10.31, p=0.0148), but no main effects of training schedule ($F_{(1,7)}$=0.3980, p=0.5481; $F_{(1,7)}$=0.002485, p=0.9616), cue extinction ($F_{(1,7)}$=0.6542, p=0.4452; $F_{(1,7)}$=1.620, p=0.2437), and there were no cue reinforcement × training schedule ($F_{(1,7)}$=3.054, p=0.1241; $F_{(1,7)}$=0.4782, p=0.5115), cue reinforcement × cue extinction ($F_{(1,7)}$=2.205, p=0.1812; $F_{(1,7)}$=1.304, p=0.2910), training schedule × cue extinction ($F_{(1,7)}$=0.03134, p=0.8645; $F_{(1,7)}$=0.001651, p=0.9687), or 3-way interactions ($F_{(1,7)}$=5.232, p=0.560; $F_{(1,7)}$=0.1821, p=0.6824). Together, there was no effect of cue extinction or interaction between cue extinction and another factor on calcium or dopamine AUC in the DLS or DMS, which indicates that cue extinction may not impact AUC in the DMS despite its effects on DMS calcium and dopamine peak amplitudes (Figure 4).

**Dorsal striatal responses to noncontingent cues during cue extinction**

After 20 days of self-administration, rats underwent cue extinction, during which levers were retracted and 120 20-second audiovisual cues were passively presented every 30 seconds over one hour. Rats used in the experiments presented in the main text underwent photometry recordings during the entire 1-hour cue extinction procedure, when 120 cues were presented noncontingently, in order to determine if there was a change in dorsal striatal responses to cues as the cue extinction session progressed. There were no differences in calcium or dopamine responses between FR-trained and SO-trained rats, so groups were combined for analysis. To account for a shift in baseline that we observed throughout cue extinction sessions, event traces were normalized by subtracting the average fluorescence during the 3-second period before cue onset from each event's trace. Sessions were divided into 15-minute, 30-cue bins and analyzed with a one-way rmANOVA.

There was no main effect of bin on DLS calcium peak amplitude ($F_{(1.496,14.96)}$=3.855, p=0.0549) (Figure S5A) or AUC ($F_{(1.801,18.01)}$=3.362, p=0.0617) (Figure S5B). For DMS calcium peak amplitude, there was a main effect of bin on peak amplitude ($F_{(2.096,20.96)}$=10.16, p=0.0007) (Figure S5C) and AUC ($F_{(2.013,20.13)}$=, p=0.0012) (Figure S5D). Post-hoc analyses indicated that DMS calcium peak amplitude for cues 1-30 was significantly greater than for cues 61-90 (p=0.0007, q=8.378) and for cues 91-120 (p=0.0113, q=5.655) (Figure S5C), and DMS calcium AUC for cues 1-30 was significantly greater than for cues 61-90 (p=0.0059, q=6.255) and for cues 91-120 (p=0.0136, q=5.492) (Tukey's multiple comparisons) (Figure S5D). There was no main effect of bin on DLS dopamine peak amplitude ($F_{(2.299,22.99)}$=1.564, p=0.2295) (Figure S5E), DLS dopamine AUC ($F_{(2.154,21.54)}$=2.960, p=0.0698) (Figure S5F), DMS dopamine peak amplitude ($F_{(1.936,19.36)}$=1.581, p=0.2313) (Figure S5G), or DMS dopamine AUC ($F_{(1.642,16.42)}$=1.245, p=0.3060) (Figure S5H).

Although these results suggested that DMS calcium responses to noncontingent cues decreased throughout the hour-long cue extinction session, we were concerned that the reduction in calcium signal could be due to photobleaching of the calcium sensor, which can occur when fluorescent sensors are excited over longer periods of time. Therefore, we decided to perform a control experiment in a separate set of animals (n=7, 2 males and 5 females, with 2 DMS fibers excluded due to fiber misplacement). These animals were trained to self-administer cocaine for 20 days (10 days on FR schedules of reinforcement followed by 10 days on SO schedules of reinforcement, similarly to SO-trained rats used throughout the main text). Rats in this control experiment also underwent cue extinction, when 120 20-second cues were presented noncontingently, but photometry recordings only occurred during cues 1-30 (first 15 minutes) and cues 91-120 (last 15 minutes), and rats were disconnected from the laser and optic cables during cues 31-90 (middle 30 minutes) to minimize potential photobleaching. Dorsal striatal responses were compared between the first and last 30 cues within subjects using paired *t*-tests, and there were no differences in DLS calcium peak z-score amplitude (p=0.3011, $\eta^2$=0.1758) (Figure S5I), DLS calcium AUC (p=0.9742, $\eta^2$=0.0003215) (Figure S5J), DMS calcium peak (p=0.6803, $\eta^2$=0.04688) (Figure S5K), DMS calcium AUC (p=0.9192, $\eta^2$=0.002907) (Figure S5L), DLS dopamine peak (p=0.8801, $\eta^2$=0.004108) (Figure S5M), or DLS dopamine AUC (p=0.6987, $\eta^2$=0.02677) (Figure S5N). There was a significant reduction in DMS dopamine peak amplitude (p=0.0062, $\eta^2$=0.8740) (Figure S5O) and AUC (p=0.0011, $\eta^2$=0.9456) (Figure S5P) during the last 30 cues. Because the initial DMS dopamine response to noncontingent cues was minimal, this reduction appeared to be due to a reduction in DMS dopamine, below baseline, upon cue onset during the last 30 cues. The results from this control experiment suggest that the reduction in DMS calcium that occurred throughout cue extinction in the rats used in the main text was likely due to photobleaching of the fluorescent sensor. However, the control experiment suggests that DMS dopamine responses to noncontingent cues may be impacted throughout cue extinction.

**Prior to operant behavioral training, novel stimulus presentation induces increases in dorsal striatal calcium and dopamine activity**

In a subset of rats (n=5), we evaluated if novel stimuli presented noncontingently prior to operant behavioral training impacted dorsal striatal calcium or dopamine activity. Rats underwent 2 days of 15-minute testing sessions in which novel stimuli were presented for 10 seconds (houselight, cue light, audio tone, levers inserted, or simultaneous cue light and audio tone) 4 times each. The z-score for each trace for each event was averaged for each rat, and the peak amplitude and average AUC per second were compared between the 3 seconds before stimulus onset (baseline) and 10 seconds during stimulus presentation using 2-way ANOVAs. For DLS calcium peak amplitude, there was a main effect of stimulus presentation ($F_{(1,4)}$=439.3, p<0.0001), but no main effect of stimulus type ($F_{(2.311,9.245)}$=3.013, p=0.0937) or stimulus presentation × stimulus type interaction ($F_{(1.723,6.891)}$=1.454, p=0.2923) (Figure S6A). There was a significant stimulus presentation × stimulus type interaction ($F_{(1.647,6.588)}$=9.868, p=0.0121) for DLS calcium AUC, and post-hoc analyses revealed a significant effect of simultaneous cue light and tone presentation (p=0.0014, t=12.06) (Sidak's multiple comparisons) (Figure S6B). For DMS calcium peak amplitude, there was a main effect of stimulus presentation ($F_{(1,4)}$=78.51, p=0.0009), but no main effect of stimulus type ($F_{(1.382,5.311)}$=2.636, p=0.1626) or stimulus presentation × stimulus type interaction ($F_{(1.291,5.163)}$=2.515, p=0.1735) (Figure S6C). For DMS calcium AUC, there was a significant stimulus presentation × stimulus type interaction ($F_{(2.383,9.527)}$=7.638, p=0.0087), but post-hoc analyses did not reveal any additional effects (Sidak's multiple comparisons) (Figure S6D). For both DLS (Figure S6E) and DMS (Figure S6G) dopamine peak amplitude, there was a main effect of stimulus presentation ($F_{(1,4)}$=51.27, p=0.0020; $F_{(1,4)}$=19.08, p=0.0120), but no main effect of stimulus type ($F_{(2.156,8.623)}$=3.002, p=0.1003; $F_{(1.950,7.798)}$=1.990, p=0.2005) or stimulus presentation × stimulus type interaction ($F_{(1.369,5.477)}$=1.137, p=0.3577; $F_{(2.271,9.084)}$=0.4416, p=0.6796). For both DLS (Figure S6F) and DMS (Figure S6H) dopamine AUC, there were no main effects of stimulus presentation ($F_{(1,4)}$=0.4533, p=0.5377; $F_{(1,4)}$=0.4211, p=0.5517) or stimulus type ($F_{(2.425,9.698)}$=2.449, p=0.1323; $F_{(1.311,5.234)}$=0.2945, p=0.6694), nor and there was no stimulus presentation × stimulus type interaction ($F_{(2.277,9.109)}$=2.236, p=0.1594; $F_{(1.302,5.208)}$=0.2718, p=0.6828). Interestingly, these data indicate that novel stimulus presentation may induce some dorsal striatal calcium and dopamine activity.

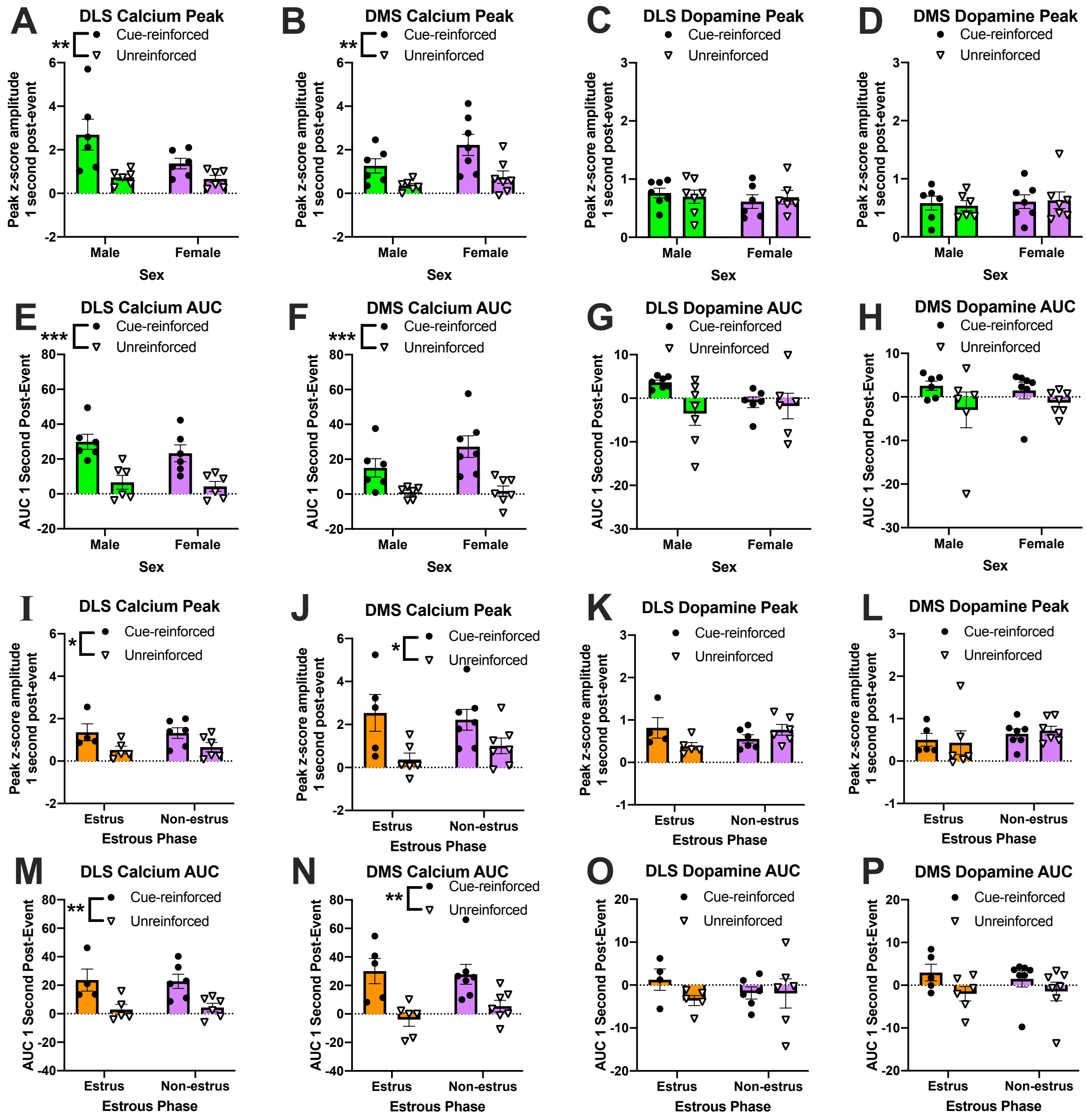

**Fig. S1. No effect of sex or estrous phase on dorsal striatal calcium or dopamine responses to lever presses.**

Because male and female rats were used for these experiments, we evaluated if sex (**A-H**) or estrous phase in females (**I-P**) had an effect on dorsal striatal calcium and dopamine responses to lever presses during the early phase of training, because we could collapse across future training schedule. For DLS calcium peak (**A**) and AUC (**E**) and DMS calcium peak (**B**) and AUC (**F**), there was a main effect of cue reinforcement but no effect of sex or cue reinforcement × sex interaction. For DLS dopamine peak (**C**) and AUC (**G**) and DMS dopamine peak (**D**) and AUC (**H**), there was no effect of cue reinforcement, sex, or interaction. For DLS calcium peak (**I**) and AUC (**M**) and DMS calcium peak (**J**) and AUC (**N**), there was a main effect of cue reinforcement but no effect of estrous phase or cue reinforcement × estrous phase interaction. For DLS dopamine peak (**K**) and AUC (**O**) and DMS dopamine peak (**L**) and AUC (**P**), there was no effect of cue reinforcement, estrous phase, or interaction. Graphs show group means ± SEM and individual data points. *$p<0.05$; **$p<0.01$; ***$p<0.001$.

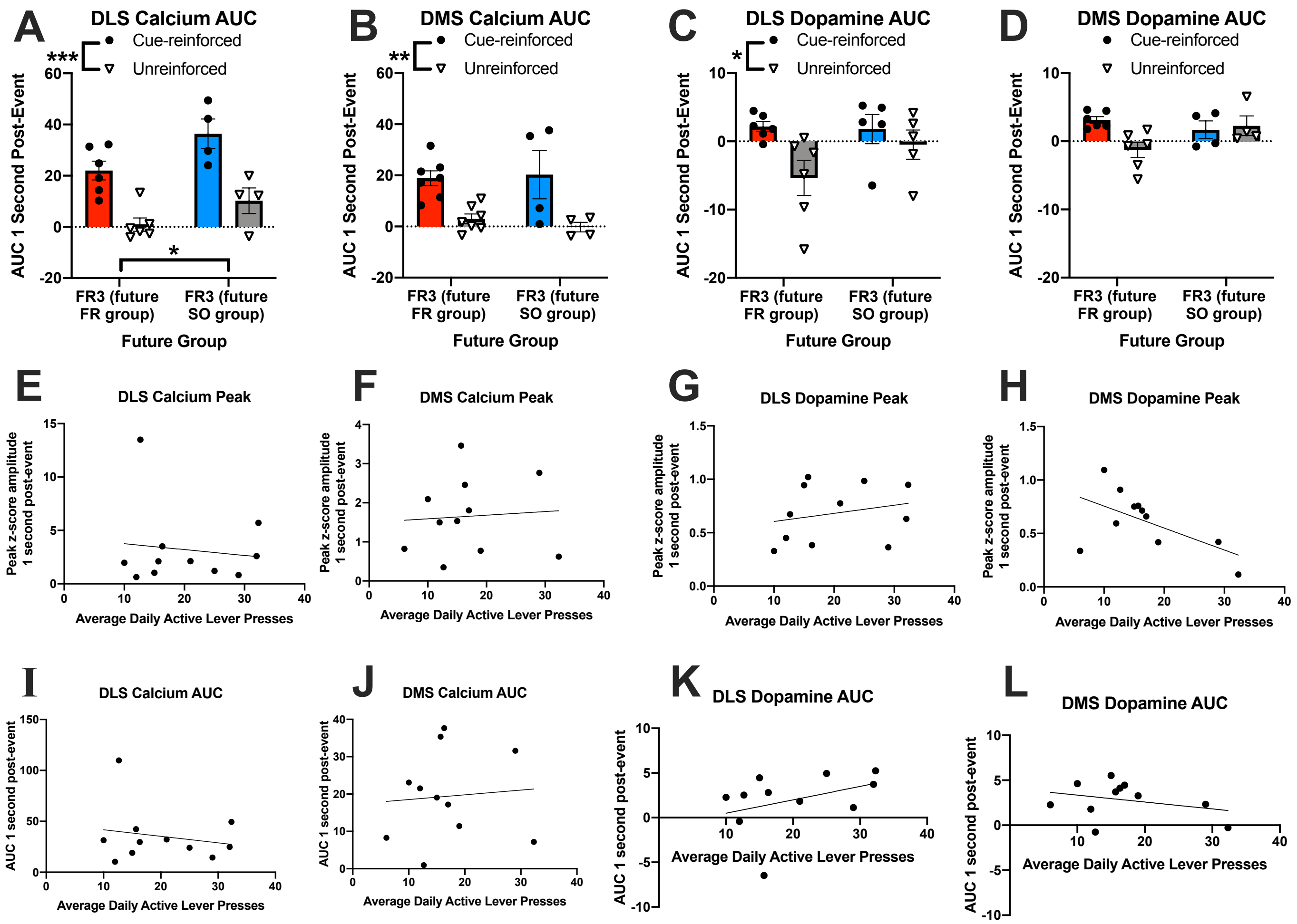


**Fig. S2. During early training, cue-reinforced lever presses resulted in greater AUC for DLS calcium and dopamine and DMS calcium, and there were no correlations between average daily active lever presses and calcium or dopamine responses to cue-reinforced lever presses.**

Initial fiber photometry recordings took place during the early phase of training, when all rats were on an FR3 schedule and had not yet been split into FR- and SO-trained groups. During this early phase of training, there was a main effect of cue reinforcement and future training schedule on DLS calcium AUC in the 1 second after lever press, but no cue reinforcement × future training schedule interaction (**A**). For DMS calcium AUC, there was a main effect of cue reinforcement, but no effect of future training schedule or interaction (**B**). There was a main effect of cue reinforcement on DLS dopamine AUC, but no main effect of future training schedule or interaction (**C**). There was no main effect of cue reinforcement or future training schedule or interaction for DMS dopamine AUC (**D**). There was no significant correlation between average daily active lever presses during early training and DLS calcium (**E**), DMS calcium (**F**), DLS dopamine

(**G**), or DMS dopamine (**H**) peak amplitudes or with DLS calcium (**I**), DMS calcium (**J**), DLS dopamine (**K**), or DMS dopamine (**L**) AUC. Graphs show group means ± SEM and individual data points. *p<0.05; **p<0.01; ***p<0.001.

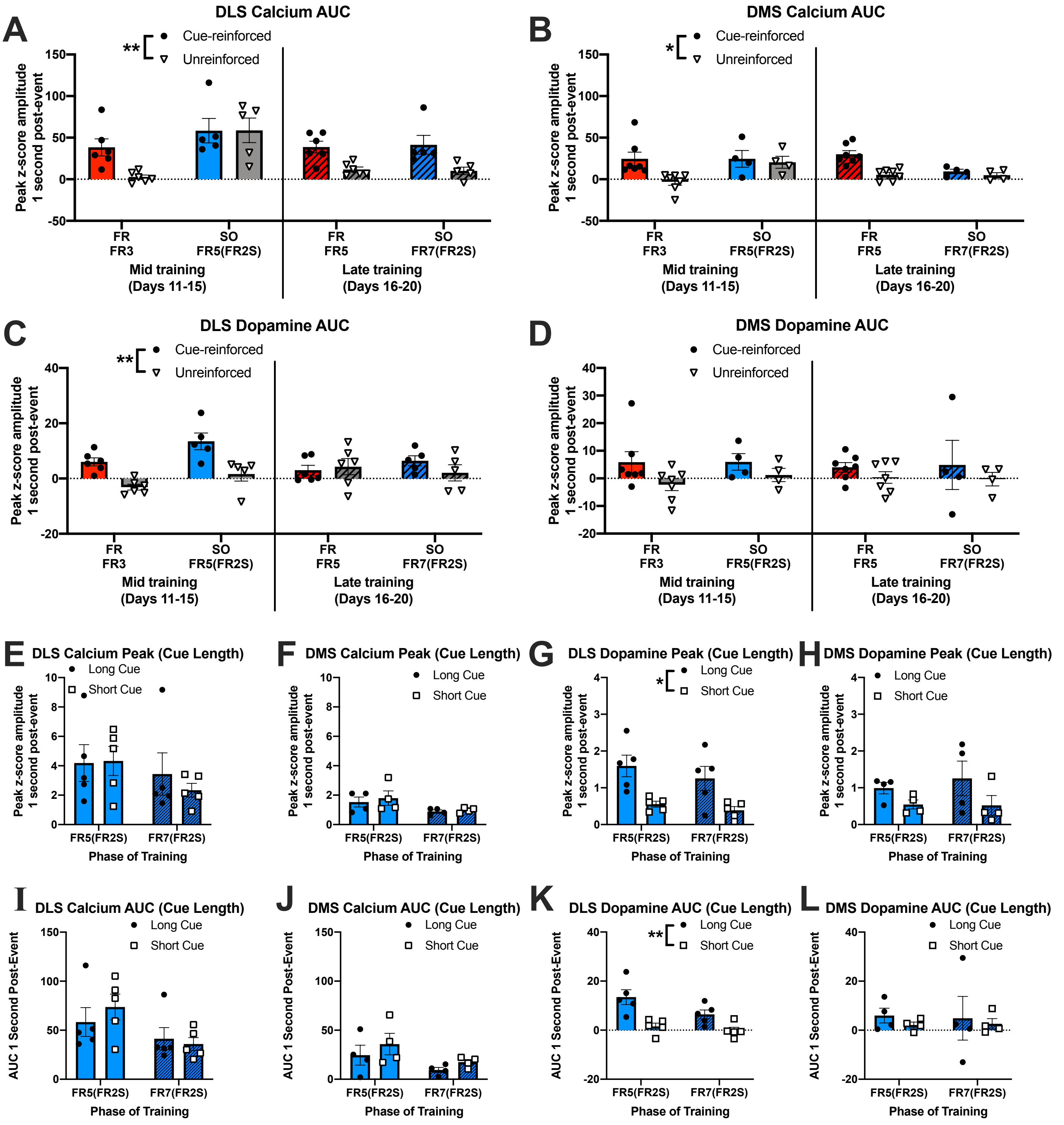


**Fig. S3. Dorsal striatal calcium and dopamine AUC after cue-reinforced lever presses differ between FR-trained and SO-trained rats, and SO-trained rats have different DLS dopamine responses to short cues vs long cues.**

During the middle and late phases of training, AUC in the 1 second after cue-reinforced and unreinforced lever presses were compared between groups. There was a main effect of cue reinforcement and training schedule on DLS calcium

AUC as well as a training schedule × phase of training interaction, but there was no main effect of phase of training or other interactions (**A**). There was a main effect of cue reinforcement on DMS calcium AUC as well as a training schedule × phase of training interaction, but there was no main effect of training schedule or phase of training or other significant interactions (**B**). For DLS dopamine AUC, there was a main effect of cue reinforcement as well as cue reinforcement × phase of training and training schedule × phase of training interactions, but there were no other main effects or interactions (**C**). For DMS dopamine AUC, there were no main effects of cue reinforcement, training schedule, or phase of training, nor were there any interactions (**D**). During SO training, rats received both short, 1-second cues upon completion of the first-order schedule and long, 20-second cues accompanied by timeout upon completion of the second-order schedule. Therefore, we compared dorsal striatal calcium and dopamine responses to lever presses that resulted in short or long cue presentation in SO-trained rats. There was no main effect of phase of training, cue length, or phase of training × cue length interaction for DLS (**E**) or DMS (**F**) calcium peak amplitude. There was a main effect of cue length on DLS dopamine peak amplitude (**G**), but no effect of phase of training or interaction. There were no main effects of phase of training, cue length, or phase of training × cue length interaction for DMS dopamine peak amplitude (**H**), DLS calcium AUC (**I**), or DMS calcium AUC (**J**). There was a main effect of cue length, but no effect of phase of training or interaction, for DLS dopamine AUC (**K**), but there were no main effects or interactions for DMS dopamine AUC (**L**). Graphs show group means ± SEM and individual data points. *$p<0.05$; **$p<0.01$.

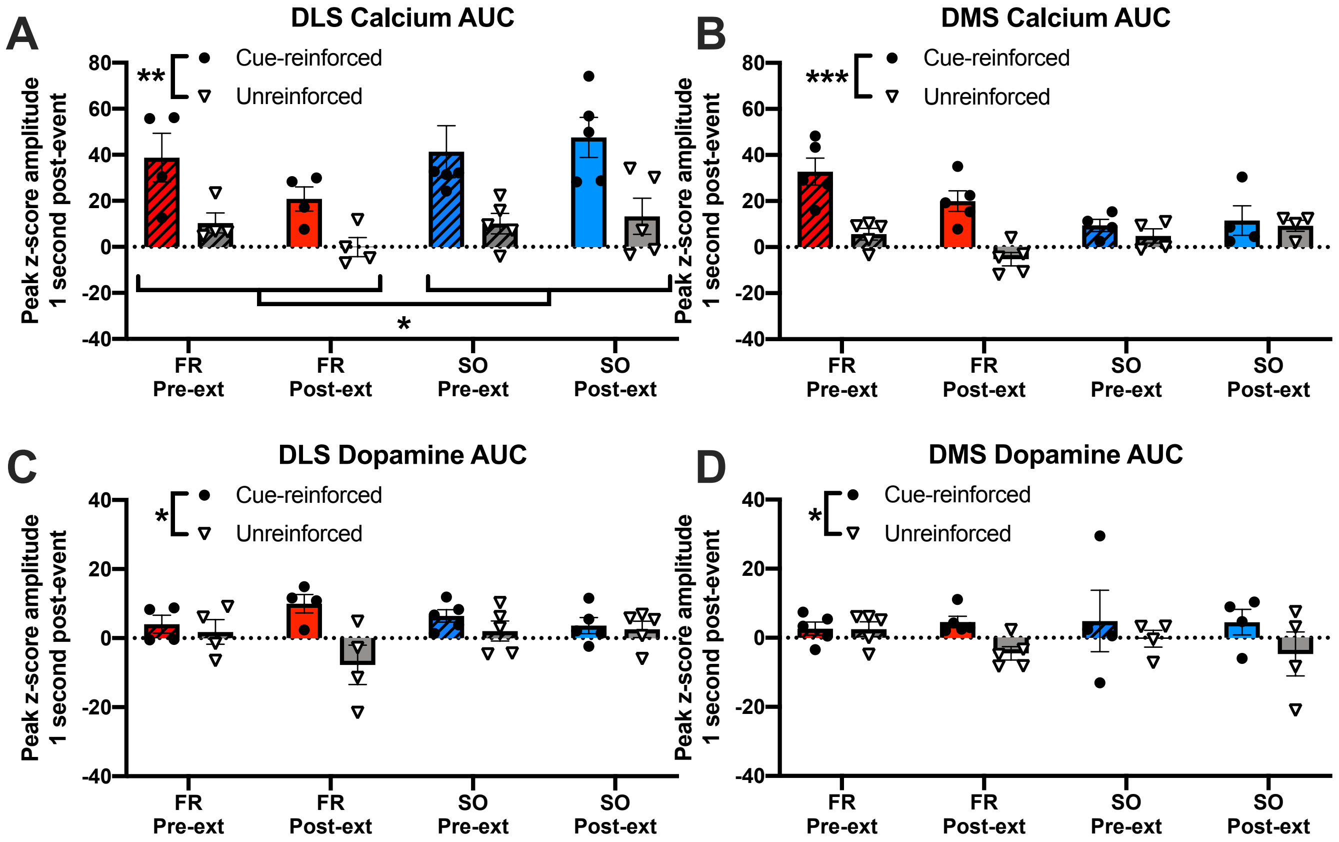


**Fig. S4. Cue extinction does not impact dorsal striatal calcium or dopamine AUC responses to lever presses.**

Dorsal striatal calcium and dopamine AUC after lever presses were compared between drug-seeking tests during the late phase of training (pre-ext) and after cue extinction (post-ext). There was a main effect of cue reinforcement and training schedule on DLS calcium AUC, but there was no main effect of cue extinction or cue reinforcement × training schedule, cue reinforcement × cue extinction, training schedule × cue extinction, or 3-way interaction (A). There was a main effect of cue reinforcement and a cue reinforcement × training schedule interaction for DMS calcium AUC, but no other main effects or interactions (B). There was a main effect of cue reinforcement on DLS dopamine AUC (C) and DMS dopamine AUC (D), but no other main effects or interactions. Graphs show group means ± SEM and individual data points. *$p<0.05$; **$p<0.01$; ***$p<0.001$.

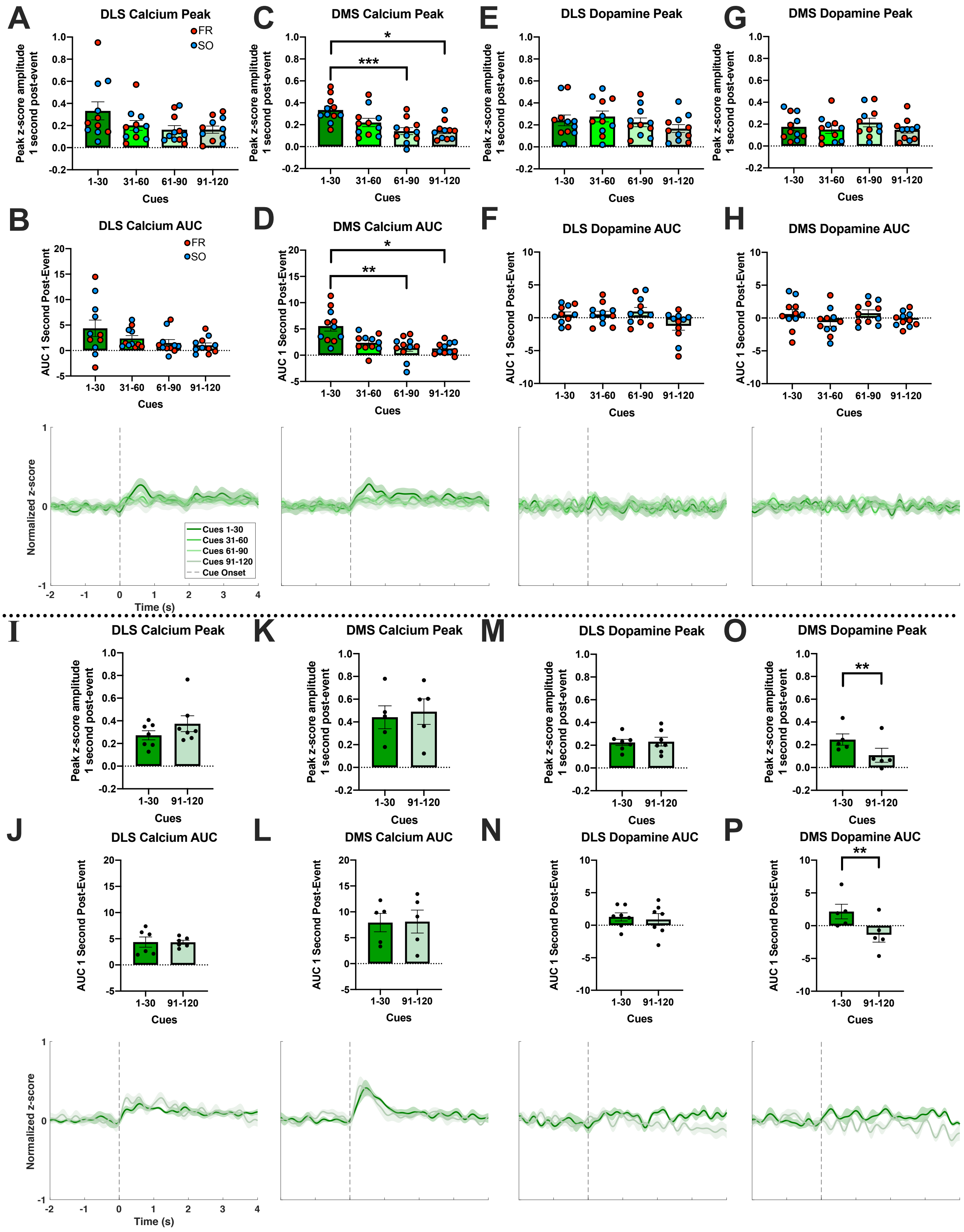
A
DLS Calcium Peak
FR
SO
Peak z-score amplitude 1 second post-event
Cues
1-30
31-60
61-90
91-120
B
DLS Calcium AUC
AUC 1 Second Post-Event
C
DMS Calcium Peak
*
***
D
DMS Calcium AUC
*
**
E
DLS Dopamine Peak
F
DLS Dopamine AUC
G
DMS Dopamine Peak
H
DMS Dopamine AUC
Normalized z-score
Time (s)
Cues 1-30
Cues 31-60
Cues 61-90
Cues 91-120
Cue Onset
I
DLS Calcium Peak
J
DLS Calcium AUC
K
DMS Calcium Peak
L
DMS Calcium AUC
M
DLS Dopamine Peak
N
DLS Dopamine AUC
O
DMS Dopamine Peak
**
P
DMS Dopamine AUC
**

**Fig. S5. Dorsal striatal responses to noncontingent cues during cue extinction.**

Dorsal striatal calcium and dopamine activity were recorded during cue extinction, when 120 20-second audiovisual cues were passively presented every 30 seconds over one hour. In the rats used throughout the main text, photometry recordings occurred throughout the 1-hour cue extinction session, and cues were separated into 15-minute, 30-cue bins. Data from FR-trained and SO-trained rats were combined for analysis. There was no main effect of bin on DLS calcium peak amplitude (**A**) or AUC (**B**). For DMS calcium peak amplitude (**C**) and AUC (**D**), there was a main effect of bin, and post-hoc analyses showed that the calcium amplitude and AUC for cues 1-30 was greater than for that of cues 61-90 and cues 91-120 (**C, D**). There was no main effect of bin on DLS dopamine peak amplitude (**E**) or AUC (**F**) or on DMS dopamine peak amplitude (**G**) or AUC (**H**). In a control experiment using different rats (n=7) trained similarly to SO-trained rats used in the main text, photometry recordings only took place during the first and last 30 cues to minimize photobleaching that could occur during long recording periods. In this experiment, there was change in DLS calcium peak amplitude (**I**) or AUC (**J**), DMS calcium peak amplitude (**K**) or AUC (**L**), or DLS dopamine peak amplitude (**M**) or AUC (**N**) between responses to the first and last 30 cues. There was a significant reduction in DMS dopamine peak amplitude (O) and AUC (P) during the last 30 cues of cue extinction. Graphs show group means ± SEM and individual data points, with FR-trained rats indicated by red symbols and SO-trained rats indicated by blue symbols. Different rats were used for experiments above and below the dashed line. Traces show overall average trace for each bin aligned to cue onset with SEM shown with shading and dashed vertical lines indicating time of cue onset. *p<0.05; **p<0.01; ***p<0.001.

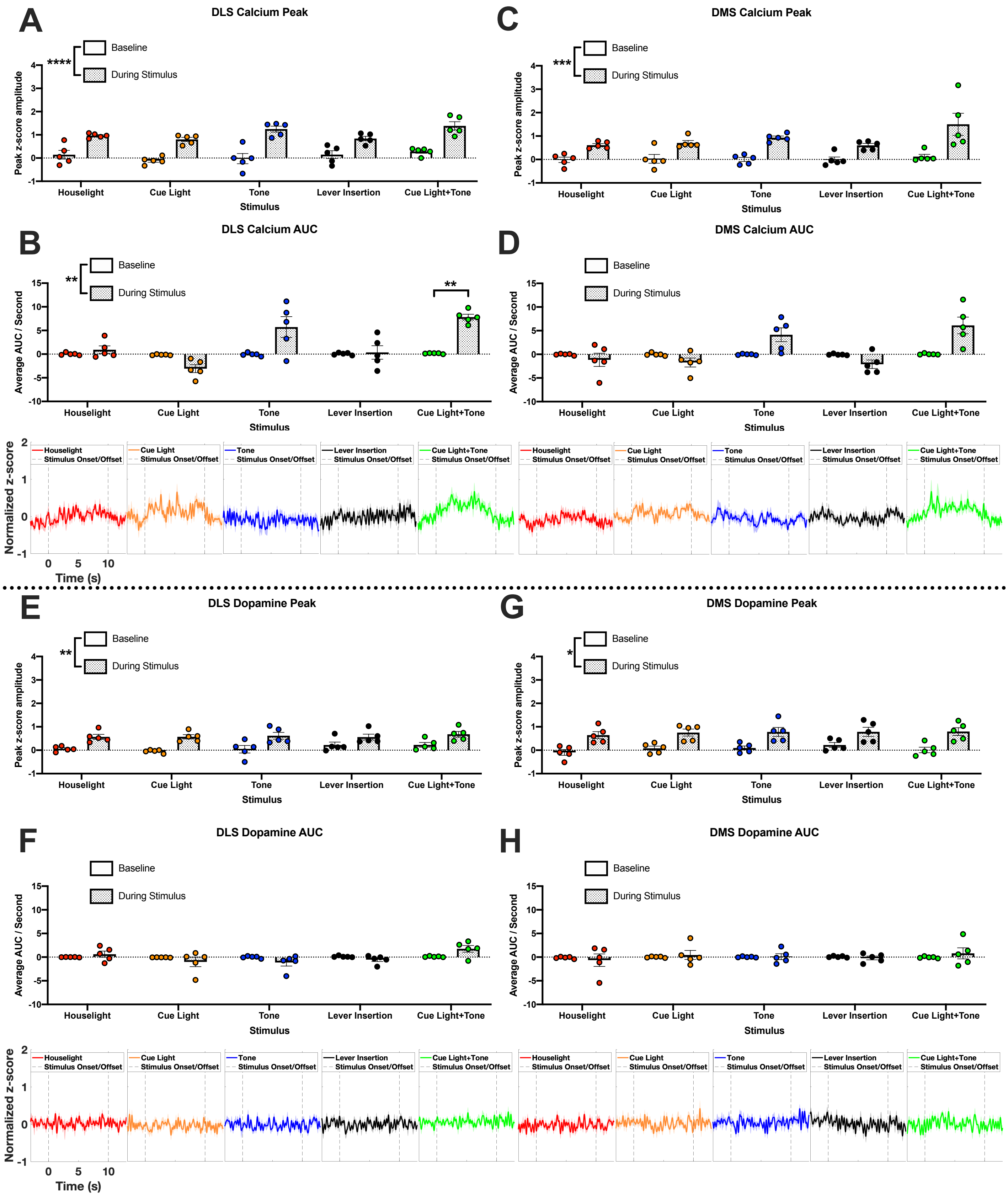


**Fig. S6. Prior to operant behavioral training, novel stimulus presentation induces increases in dorsal striatal**

**calcium and dopamine activity.**

A subset of rats (n=5) were exposed to novel stimuli (houselight, cue light, audio tone, levers inserted, or simultaneous cue light and audio tone) prior to operant behavioral training. The effect of these stimuli on dorsal striatal peak amplitude and average AUC per second were compared between the 3 seconds before stimulus onset and 10 seconds during stimulus onset. For DLS calcium peak amplitude, there was a main effect of stimulus presentation, but no main effect of stimulus type or stimulus presentation × stimulus type interaction (**A**). For DLS calcium AUC, there was a significant stimulus presentation × stimulus type interaction, and post-hoc analyses revealed a significant effect of simultaneous cue light and tone presentation on DLS calcium AUC (**B**). There was a main effect of stimulus presentation on DMS calcium peak amplitude, but no main effect of stimulus type or stimulus presentation × stimulus type interaction (**C**). For DMS calcium AUC, there was a significant stimulus presentation × stimulus type interaction, but post-hoc analyses did not reveal any additional effects (**D**). There was a main effect of stimulus presentation, but no main effect of stimulus type or interaction, on DLS dopamine peak amplitude (**E**), but there were no main effects or interactions for DLS dopamine AUC (**F**). Similarly, there was a main effect of stimulus presentation, but no main effect of stimulus type or interaction, on DMS dopamine peak amplitude (**G**), but there were no main effects or interactions for DMS dopamine AUC (**H**). Graphs show group means ± SEM and individual data points. Traces show overall average trace for each event aligned to behavioral events with SEM shown with shading and dashed vertical lines indicating stimulus onset and offset. *$p<0.05$; **$p<0.01$; ***$p<0.001$; ****$p<0.0001$.
